# Evolution towards simplicity in bacterial small heat shock protein system

**DOI:** 10.1101/2023.05.18.541272

**Authors:** Piotr Karaś, Klaudia Kochanowicz, Marcin Pitek, Przemyslaw Domanski, Igor Obuchowski, Bartlomiej Tomiczek, Krzysztof Liberek

**Author notes:** Correspondence to B.T. or K.L.

## Abstract

Evolution can tinker with multi-protein machines and replace them with simpler single-protein systems performing equivalent functions in equally efficient manner. It is unclear how, on a molecular level, such simplification can arise. With ancestral reconstruction and biochemical analysis we have traced the evolution of bacterial small heat shock proteins (sHsp), which help to refold proteins from aggregates using either two proteins with different functions (IbpA and IbpB) or a secondarily single sHsp that performs both functions in an equally efficient way. Secondarily single sHsp evolved from IbpA, an ancestor specialized in strong substrate binding. Evolution of an intermolecular binding site drove the alteration of substrate binding properties, as well as formation of higher-order oligomers. Upon two mutations in the α-crystallin domain, secondarily single sHsp interacts with aggregated substrates less tightly. Paradoxically, less efficient binding positively influences the ability of sHsp to stimulate substrate refolding, since the dissociation of sHps from aggregates is required to initiate Hsp70-Hsp100-dependent substrate refolding. After the loss of a partner, IbpA took over its role in facilitating the sHsp dissociation from an aggregate by weakening the interaction with the substrate, which became beneficial for the refolding process. We show that the same two amino acids introduced in modern-day system define whether the IbpA acts as a single sHsp or obligatorily cooperates with an IbpB partner. Our discoveries illuminate how one sequence has evolved to encode functions previously performed by two distinct proteins.

## Introduction

Gene birth and loss is a hallmark of protein family evolution, however molecular determinants and genetic mechanisms enabling that process are not well understood (Fernandez & Gabaldon, 2020; Worth *et al*, 2009). Gene loss and differential retention of paralogues reshapes the divergence of organisms both in *Animalia, Archaea* and *Bacteria* (Fernandez & Gabaldon, 2020; Iranzo *et al*, 2019; Puigbo *et al*, 2014). Examples of gene loss often involve adaptive changes in response to changing environmental niches, like loss of genes encoding olfactory receptors in primates (Demuth & Hahn, 2009) or differential retention of paralogous genes encoding venom toxins in different rattlesnake lineages (Dowell *et al*, 2016). In prokaryote genomes, gene loss is one of the main evolutionary processes accelerating sequence divergence leading to functional innovations (Puigbo *et al*., 2014). In complex protein systems execution of a cellular function can be shared between several proteins. In bacteria the cost of maintaining additional gene copy is very high and maintaining low gene count is important for keeping the replication energy costs low (Kempes *et al*, 2017; Lever *et al*, 2015; Lynch & Marinov, 2015). Still, it remains unclear how a multi-protein system can undergo simplification. Here we asked what are the molecular events that enabled the gene loss, and how one of the biochemical functions has been taken over by the other protein. We investigated these questions using small heat shock protein (sHsp) system, which underwent simplification within *Enterobacterales* (which include common bacteria species like *Escherichia coli, Salmonella enterica* and *Erwinia amylovora)* as a model (Fig. 1).

**Fig. 1.**
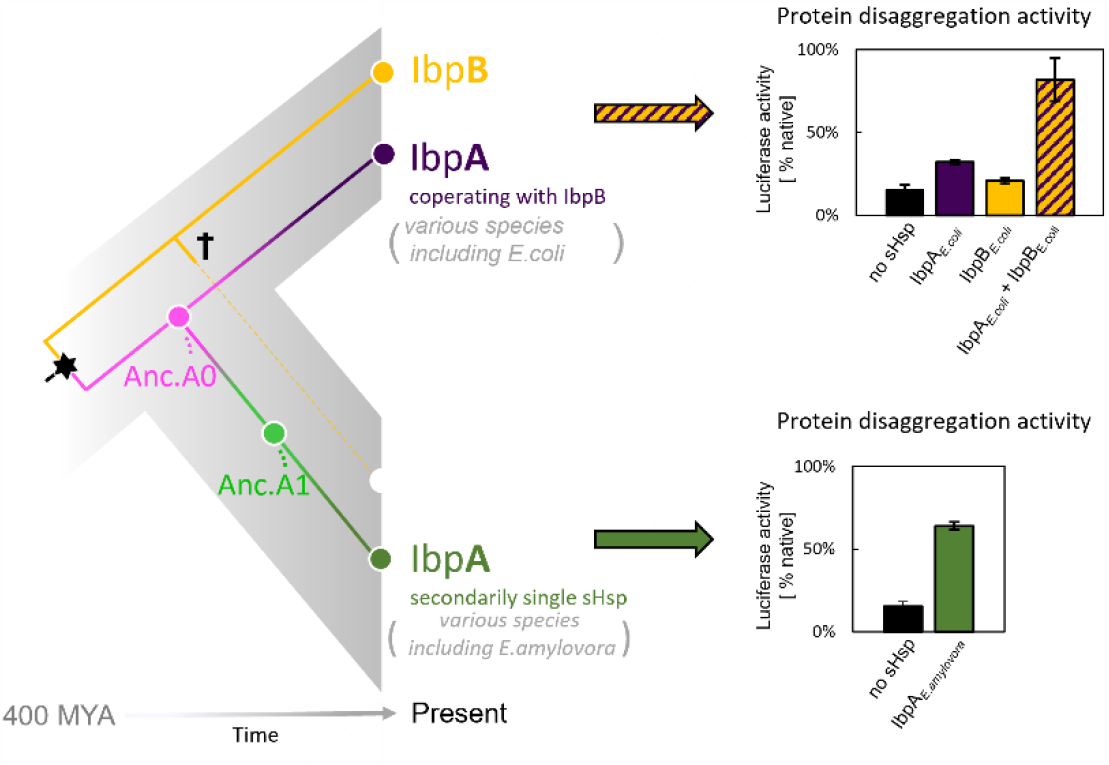
sHsp systems in *Enterobacteriacrae* and *Erwiniaceae*. Left - schematic phylogeny of sHsps in *Enterobacterales*. Gene duplication resulting in IbpA + IbpB two-protein system is marked with a star, while the loss of *ibpB* gene in *Erwiniaceae* clade is marked with a cross. AncA_0_ – reconstructed last common ancestor of IbpA from *Erwiniaceae* and *Enterobacteriaceae*, expressed as a part of two-protein system. AncA_1_ – reconstructed last common ancestor of secondarily single IbpA from *Erwiniaceae*. Right - representative extant sHsps’ ability to stimulate luciferase refolding. sHsps were present during the luciferase thermal denaturation step. Refolding of denatured luciferase was performed by the Hsp70-Hsp100 chaperone system. Activity of luciferase was measured after 1h refolding at 25 °C and shown as an average of at least three repeats ± standard deviation.

sHsps are a family of ATP – independent molecular chaperones present in all living organisms with various copy numbers (ten representatives in human) (Haslbeck & Vierling, 2015). They bind misfolded proteins and sequester them into refolding – prone assemblies, preventing uncontrolled aggregation and helping to maintain proteostasis at stress conditions. sHsp are composed of a highly conserved α–crystallin domain (ACD), in a form of so-called β–sandwich, flanked by less conserved, unstructured N – and C – terminal regions (Haslbeck & Vierling, 2015; Haslbeck *et al*, 2019; Reinle *et al*, 2022). Their smallest functional unit is usually a dimer, formed by the interaction between ACDs of two neighboring sHsps. Stable sHsp dimers in turn tend to form variable and dynamic higher – order oligomers, stabilized by N – and C – terminal region interactions. Particularly the interaction between IXI motif, highly conserved in sHsps C-termini, and the cleft formed by β4 and β8 strands of ACD is critical for oligomer formation (Haslbeck & Vierling, 2015; Kennaway *et al*, 2005; Mani *et al*, 2016; Strozecka *et al*, 2012). Oligomers of bacterial sHsps reversibly dissociate into smaller forms when the temperature increases. It is considered their activation mechanism, probably uncovering substrate interaction sites. The mechanism of sHsps’ interaction with misfolded substrates is not yet fully understood, but both N - termini and β4 – β8 cleft region have been found to play a role in this process (Basha *et al*, 2006; Fuchs *et al*, 2009; Jaya *et al*, 2009; Lee *et al*, 1997; Reinle *et al*., 2022).

In most *Enterobacterales* a two-protein sHsps system exists, consisting of IbpA and IbpB proteins (Mogk *et al*, 2003; Obuchowski *et al*, 2019). IbpA and IbpB have originated via duplication, form a heterodimer partnership and are functionally divergent from one another (Obuchowski *et al*., 2019; Pirog *et al*, 2021) (Fig. 1A,B). IbpA is specialized in tight substrate binding (sequestrase activity), while IbpB is required for dissociation of both sHps from the aggregates, a step necessary to initiate Hsp70-Hsp100 dependent substrate disaggregation and refolding (Obuchowski *et al*, 2021; Ratajczak *et al*, 2009). In a subset of *Enterobacterales* (*Erwiniaceae*), as a result of *ibpB* gene loss, the secondarily single-protein sHsp (IbpA) system has emerged (Fig. 1). The term “secondarily single” is used in order to distinguish it from single-protein IbpA from clades in which the duplication did not occur (for example *Vibrionaceae*) (Obuchowski *et al*., 2019). How did IbpA evolve to become independent of its partner? In this study, using ancestral reconstruction, we identify mutations, which allowed the secondarily single IbpA to be fully functional without its partner in substrate sequestration and handover to Hsp70-Hsp100-mediated disaggregation and refolding.

## Results

### New activity of *Erwiniaceae* IbpA has evolved in parallel to *ibpB* gene loss

To better understand the evolution of sHsps after gene loss, we reconstructed the IbpA ancestors from before and after the loss of its IbpB partner. This technique uses multiple sequence alignments of modern-day proteins from different species to infer amino acid sequences of its common ancestors (Ashkenazy *et al*, 2012; Pupko *et al*, 2002) and is widely used to investigate various evolutionary questions (Gaucher *et al*, 2008; Longo *et al*, 2020; Thomson *et al*, 2005; Thornton *et al*, 2003). We created a multiple sequence alignment of 77 IbpA sequences from *Enterobacterales* (supplementary file 1), from which we inferred the phylogeny of IbpA using maximum likelihood method (Fig. 2, supplementary file 2 – phylogenetic tree in newick format). From that we inferred ancestral sequences, which have the highest probability of producing the modern-day sequences using the empirical Bayes method (Ashkenazy *et al*., 2012; Cohen & Pupko, 2011; Cohen *et al*, 2008; Pupko *et al*., 2002; Simmons & Ochoterena, 2000). Next, we resurrected (i.e., expressed and purified) the last ancestor of IbpA present before (AncA_0_) and after (AncA_1_) the differential gene loss (Figs. 2, supplementary file 3).

**Fig. 2.**
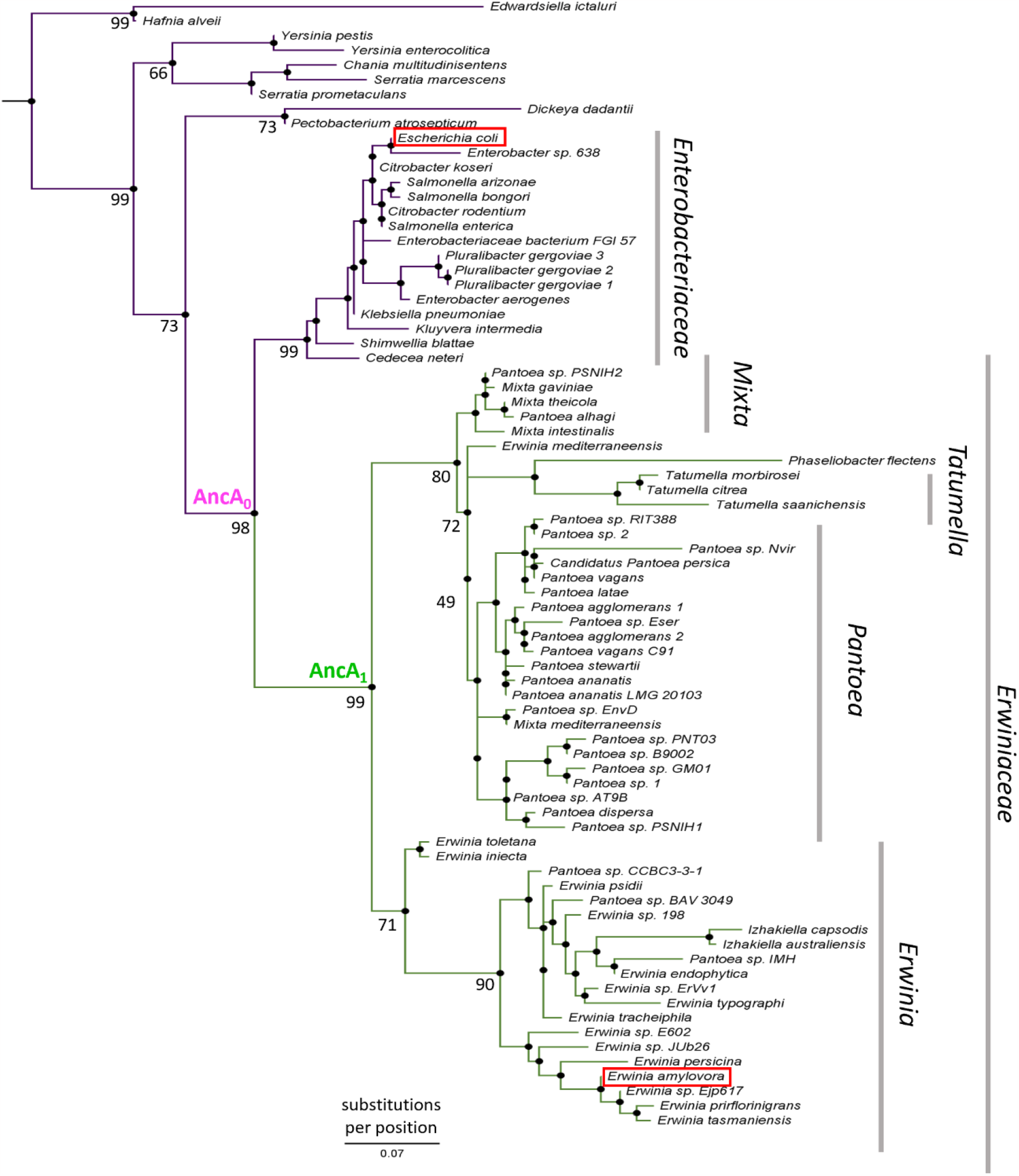
IbpA phylogeny in *Enterobacterales*: Phylogeny was reconstructed from 77 IbpA orthologs from *Enterobacterales* using Maximum Likelihood algorithm with JTT + R3 substitution model. AncA_0_ – node representing the last common ancestor of IbpA from *Erwiniaceae* and *Enterobacteriaceae*. AncA_1_ – node representing the last common ancestor of IbpA from *Erwiniaceae*. Bootstrap support is noted for the major nodes. Extant IbpAs from *E. coli* and *E. amylovora* are marked with a red frame. Scale bar – substitutions per position.

Similarly, to modern-day IbpA proteins both AncA_0_ and AncA_1_ were fully folded, and reversibly deoligomerized into smaller species under elevated temperature (Fig. 3 – figure supplement 1). Moreover, both ancestral proteins were able to sequester aggregating firefly luciferase in sHsp-substrate assemblies. AncA_0_ exhibited sequestrase activity on the level comparable to IbpA from *Escherichia coli* (IbpA_*E*.*coli*_). AncA_1_ was moderately efficient in this process and IbpA from *Erwinia amylovora* (IbpA_*E*.*amyl*_) was the least efficient sequestrase (Fig. 3A). The differences in sequestrase activity were especially pronounced at lower sHsp concentrations. Next, we tested their ability to bind protein aggregates in real time (Fig. 3B). Ancestral proteins’ interaction with the aggregated substrates was stronger than in the case of extant *E. amylovora* IbpA, but weaker than in the case of extant *E. coli* IbpA (Fig. 3B).

**Fig. 3.**
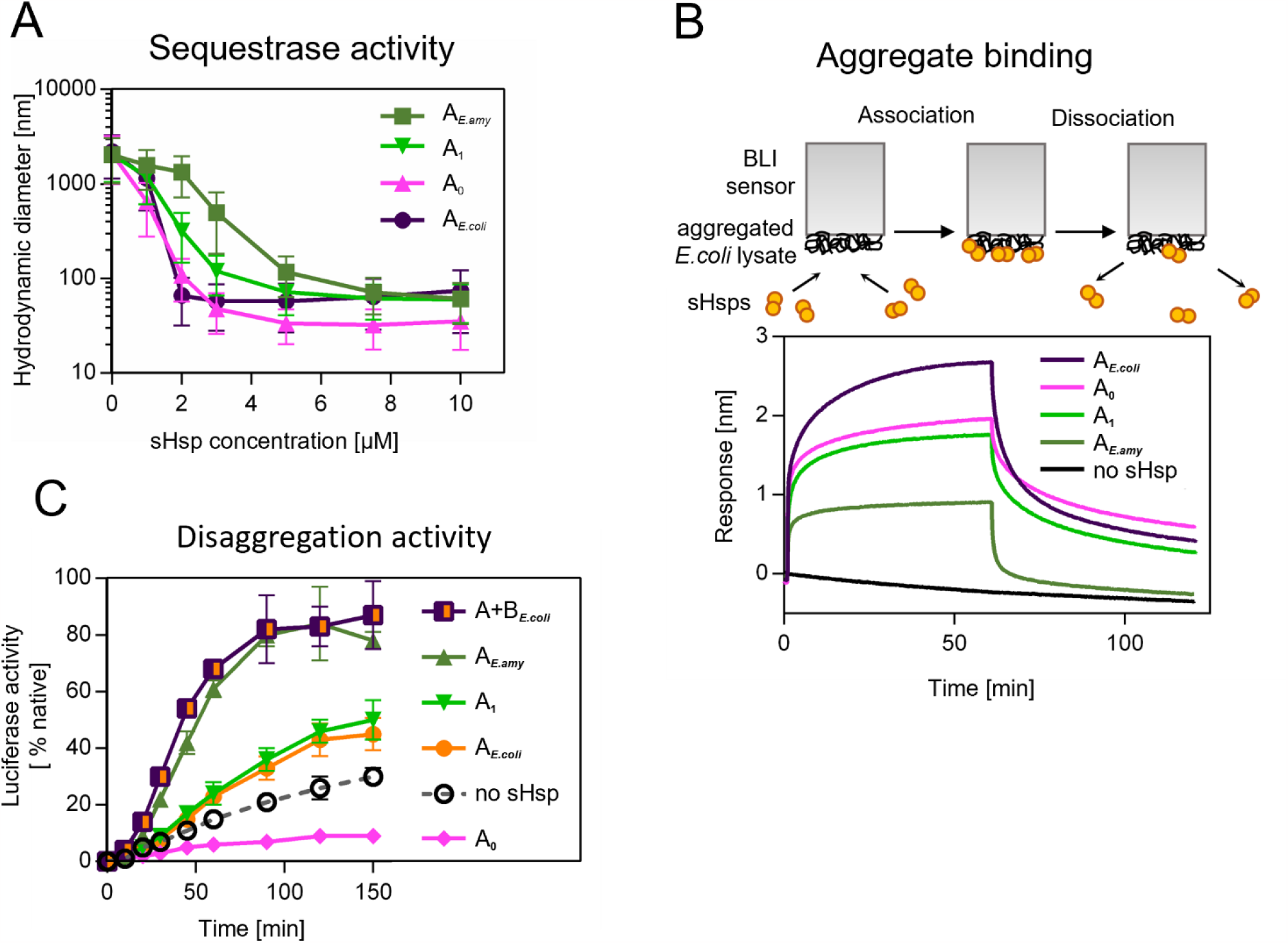
Functional changes during the evolution of secondarily single sHsp in *Erwiniaceae*. (A) Sequestrase activity of extant and ancestral sHsps. Luciferase was heat denatured in the presence of different concentrations of sHsps and size of formed sHsps – substrate assemblies was measured by DLS. Results are shown as average hydrodynamic diameter ± standard deviation. (B) Binding of extant and ancestral sHsps to heat-aggregated *E. coli* proteins. *E. coli* proteins were heat aggregated and immobilized on a BLI sensor. sHsps were heat activated before the binding step. (C) Extant and ancestral sHsps’ ability to stimulate luciferase refolding. Experiment was performed at 25 °C. Luciferase activity at each timepoint was shown as an average of at least three repeats ± standard deviation.

Finally, we asked how the modification of the substrate aggregation process by reconstructed proteins influences subsequent substrate refolding by the Hsp100 and Hsp70 chaperones. AncA_1_ stimulated luciferase refolding, however its effectiveness was around half of both analyzed extant sHsp systems (single IbpA from *E. amylovora* or IbpA + IbpB system from *E. coli*), similar to extant IbpA form *E. coli* without it’s IbpB partner. AncA_0_, in contrast, inhibited luciferase refolding in comparison to control (no sHsps at substrate aggregation step) (Fig. 3C).

To test the robustness of our observations, we repeated the analysis for the alternative ancestors, which have the second highest probability to produce the modern-day sequences (AltAll) (Fig. 3 – figure supplement 2, supplementary file 3 – posterior probabilities) (Eick *et al*, 2017). Both AltAll variants behaved similarly to most likely (ML) variants in reversible deoligomerization, sequestrase activity and stimulation of substrate refolding assays (Fig. 3 – figure supplement 3 A-D). However, the last property (the influence on refolding) required higher Hsp70 system concentration to observe AltAll variants activity (Fig. 3 – figure supplement 3 C,D). Together, these data show that reconstructed ML and AltAll ancestors are functional. What is of particular interest, these data clearly point out that the ability of reconstructed sHsps to stimulate Hsp70-Hsp100-dependent substrate refolding arose between A_0_ and A_1_ nodes.

We performed a molecular evolution analysis to test for positive selection across IbpA phylogeny using both branch models and branch-site models in codeml (Jeffares *et al*, 2015; Yang, 1998, 2007; Yang & Nielsen, 2002). The analysis shows a significantly increased ratio of nonsynonymous to synonymous substitutions, after the gene loss, at the branch leading to A_1_ with both tests. This result indicates that the new IbpA functionality likely arose due to an episode of positive selection rather than genetic drift (Fig. 4A, Supplementary file 4 A,B). The result of the branch-site test indicates possible positive selection acting on all sites substituted at the branch leading to A_1_ with pp>0.5, therefore we aimed to identify minimum number of mutations that are responsible for a change in functional properties of IbpA.

**Fig. 4.**
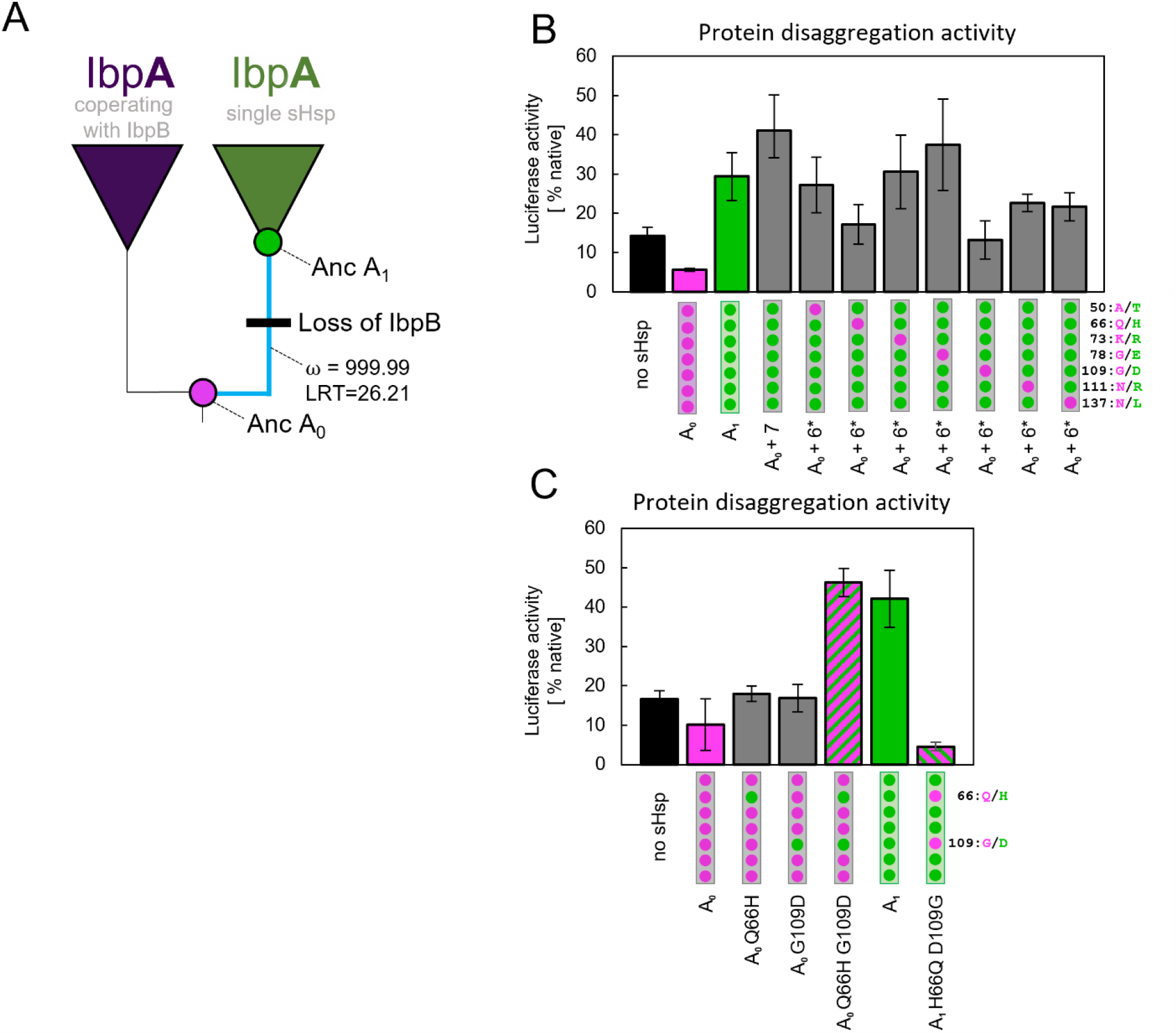
Substitutions at positions 66 and 109 that occurred between nodes A_0_ and A_1_ are crucial for ancestral sHsps to work as a single protein. Luciferase refolding assay was performed as in fig. 1. Activity of luciferase was measured after 1h refolding at 25 °C and shown as an average of at least three repeats ± standard deviation. (A) Schematic phylogeny of *Enterobacterales* IbpA showing increased ratio of nonsynonymous to synonymous substitutions (ω) on the branch between nodes AncA_0_ and AncA_1_ . Loss of cooperating IbpB is marked on a tree. Value of Likelihood Ratio Test (LRT) is given for the selection model. (B) Identification of substitutions necessary for AncA_0_ to obtain AncA_1_ – like activity in luciferase disaggregation; seven candidate mutations were introduced into AncA_0_ (AncA_0_ +7); subsequently, in series of six mutants, each of the candidate positions was reversed to a more ancestral state (AncA_0_ + 6* variants) (C) Effect of substitutions at positions 66 and 109 on the ability of AncA_0_ and AncA_1_ to stimulate luciferase refolding.

### Identification of residues defining ancestral sHsps activities

In order to identify amino acids responsible for the observed new functionality of AncA_1_, we compared the sequences of two ancestral proteins, selecting seven out of ten substitutions as probable candidates. Three substitutions were removed from analysis based on the low conservation of these positions in extant proteins (Fig. 4 – figure supplement 1). The remaining seven were introduced into AncA_0_ . Resulting AncA_0_ +7 protein stimulated Hsp70-Hsp100-dependent luciferase refolding at the level comparable to AncA_1_ (Fig. 4B). To further specify key mutations, we prepared seven additional variants. In each variant a different position in AncA_0_ +7 was reversed to a more ancestral state. The substantial decrease in luciferase refolding stimulation was observed for positions 66 and 109 (Fig. 4B). Next, each of these substitutions on its own (Q66H or G109D) was separately introduced into AncA_0_ . This was not sufficient to increase AncA_0_ ability to stimulate luciferase refolding. However, when both substitutions were introduced simultaneously, the resulting sHsp exhibited activity similar to AncA_1_ (Fig. 4C). What is more, when in AncA_1_ these two positions were reversed to AncA_0_ -like state, the resulting sHsp lost the ability to stimulate luciferase refolding (Fig. 4C). All analyzed proteins, namely AncA_0_ +7, AncA_0_ Q66H G109D and AncA_1_ H66Q D109G, possess biochemical properties characteristic for sHsps, exhibiting reversible thermal deoligomerization and sequestrase activity (Fig. 4 – figure supplement 2 A,B). All these results show that substitutions Q66H and G109D are both sufficient and necessary for the increase in activity observed for ancestral sHsps between A_0_ and A_1_ nodes.

### Identified substitutions influence α-crystallin domain properties

Substitutions Q66H and G109D, responsible for gaining single sHsp activity, are located in the α-crystallin domain (ACD) within β4 and β8 strands, which form a cleft responsible for the interaction with unstructured C-terminal peptide of the neighboring sHsp dimer (Fig. 5A). In order to identify possible structural underpinnings of the single sHsp activity we have predicted structures of AncA_0_ and AncA_0_ Q66H G109D α-crystallin domain (ACD) dimers in complex with C-terminal peptide using AlphaFold2 and *in silico* mutagenesis and subjected them to 0.5 μs equilibrium molecular dynamics (MD) simulations. Analysis of the C-terminal peptide interface contact probabilities in MD trajectories showed that both substituted residues contact the C-terminal peptide, although overall contact pattern remain similar upon their introduction (Fig. 5 – figure supplement 1) and no major differences in the overall ACD domains structure were observed (Fig. 5 – figure supplement 2) . To explore the possibility that identified substitutions affect strength of this interaction, we analyzed binding of purified ACDs of AncA_0_ and AncA_0_ Q66H G109D to C– terminal peptide using biolayer interferometry. Titrations of immobilized C-terminal peptide by different ACDs (Figs. 5B, 5 – figure supplement 3) allowed us to determine the dissociation constants. These two substitutions increased the K_0.5_ of ACD binding to the C -terminal peptide from 4.3 μM to 7.1 μM at the same time increasing the Hill coefficient of the interaction from 2.3 to 3.7, indicating a modest decrease in affinity, accompanied by an increase in binding cooperativity.

**Fig. 5.**
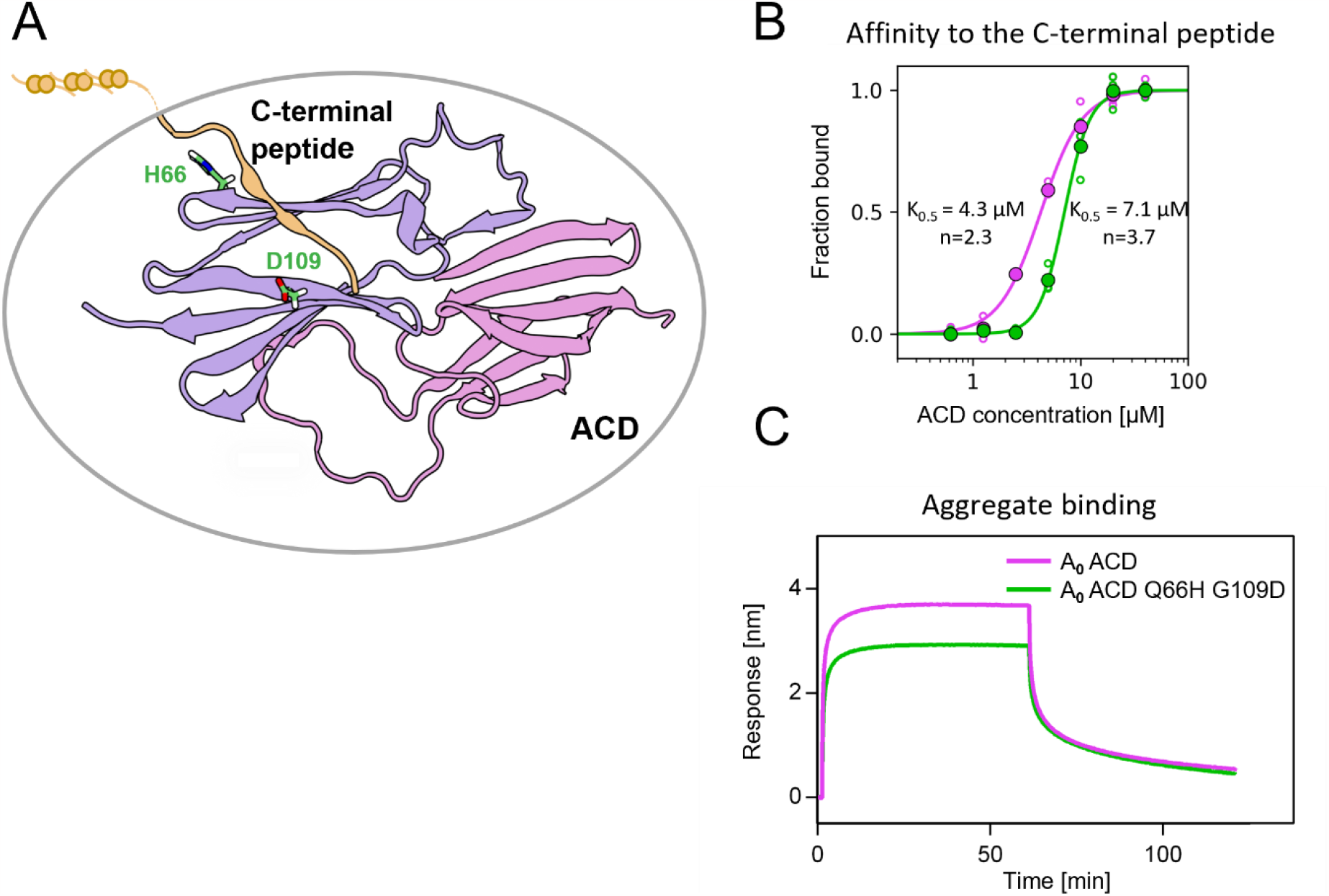
Substitutions at positions 66 and 109 decreased the affinity of AncA_0_ ACD to C-terminal peptide and aggregated substrate. (A) Structural model of complex formed by AncA_0_ Q66H G109D α-crystallin domain dimer (purple and lilac) and AncA_0_ C-terminal peptide (orange). (B) Effect of Q66H G109D substitutions (green) on AncA_0_ (purple) ACD’s affinity to the C - terminal peptide assayed by BLI. Biolayer thickness at the end of the association step was used to calculate the fraction of bound peptide. Filled circles represent means of triplicate measurements, individual data points are shown as hollow circles and were fitted to cooperative binding model (Hill equation). Values of fitted binding affinities [K_0.5_] (AncA_0_ 4.3 ±0.2 μM, AncA_0_ Q66H G109D 7.1 ±0.2 μM) and Hill coefficients [n] (AncA_0_ 2.3 ±0.17, AncA_0_ Q66H G109D 3.7 ±0.34) are indicated on the plot. (C) Effect of Q66H G109D substitutions on AncA_0_ ACD’s affinity to aggregated *E. coli* proteins bound to BLI sensor. Analysis was performed as in Fig. 3A.

As this interaction is known to play a crucial role in formation of sHsp oligomers (Fu *et al*, 2005; Mani *et al*., 2016; Strozecka *et al*., 2012), we used dynamic light scattering to investigate how Q66H and G109D substitutions influence the size of oligomers formed by AncA_0_ at different temperatures. In agreement with decreased affinity between ACD and the C–terminal peptide, we have shown that these substitutions slightly decrease the oligomer size and facilitate AncA_0_ deoligomerization (Figure 5 – figure supplement 4 A,B).

β4-β8 cleft in certain sHsps, in addition to its interaction with C – terminal peptide, was also shown to participate in interactions with substrates and partner proteins (Fuchs *et al*., 2009; Jaya *et al*., 2009; Lee *et al*., 1997; Reinle *et al*., 2022). Therefore, we tested whether ACD of AncA_0_ binds protein aggregates and whether this interaction is influenced by Q66H G109D substitutions. We observed that AncA_0_ ACD efficiently binds to either aggregated *E. coli* lysate or aggregated luciferase, and this binding was weakened by analyzed substitutions (Figs. 5C, 5 – figure supplement 5). This suggests that ACD of bacterial sHsps interacts with the substrate, most likely through the β4-β8 cleft.

These results allow us to conclude that substitutions Q66H and G109D in AncA_0_ substantially increased the sHsp ability to stimulate Hsp70-Hsp100-dependent substrate refolding by weakening the interaction of β4-β8 cleft with both the C-terminal peptide and the aggregated substrates. Despite its ability to bind aggregated substrates in biolayer interferometry assay, analyzed ACDs do not exhibit sequestrase activity and were unable to positively influence substrate refolding by the Hsp70-Hsp100 system (Fig. 5 – figure supplement 5 B,C).

### Identified substitutions define the mode of action of extant sHsps

As more ancestral, AncA_0_ -like state in positions 66 and 109 is conserved in IbpA of *E. coli* while more modern, AncA_1_ -like state is conserved in IbpA of *E. amylovora*, we decided to ask whether this difference is sufficient to explain functional differences between the two extant proteins. Therefore, we introduced AncA_1_ -like substitutions into IbpA_*E*.*coli*_ and AncA_0_ -like substitutions into IbpA_*E*.*amyl*_ . Resulting IbpA_*E*.*coli*_ Q66H G109D, in comparison to wild type IbpA_*E*.*coli*_, exhibited increased ability to stimulate Hsp70-Hsp100-dependent luciferase refolding, as well as a decreased ability to bind aggregated substrates – becoming more similar to modern *E. amylovora* IbpA. At the same time, IbpA_*E*.*amyl*_ H67Q D110G significantly less efficiently stimulated luciferase refolding, while exhibiting increased ability to bind aggregated substrates in comparison to wild type IbpA_*E*.*amyl*_ (Fig. 6A-C). Still, both new IbpA variants exhibited properties characteristic for sHsps, namely reversible thermal deoligomerization and sequestrase activity (Fig. 6 - figure supplement 1 A,B).

**Fig. 6.**
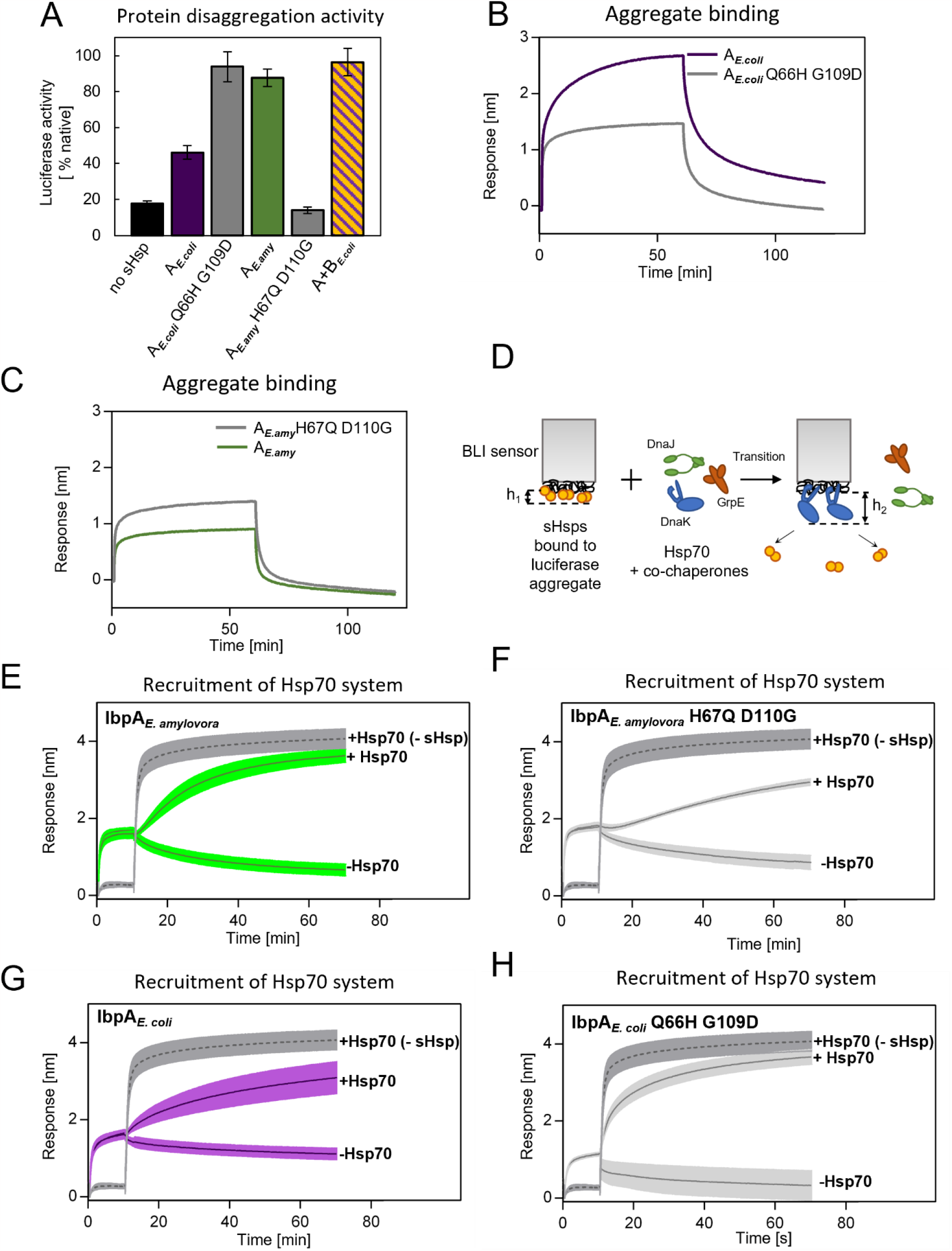
Differences at positions 66 and 109 determine functional differences between extant IbpA proteins from *E. coli* and *E. amylovora*. (A) Effect of substitutions at position 66 and 109 (and homologous) on the ability of IbpA from E. amylovora and E. coli to stimulate luciferase refolding. Assay was performed as in Fig. 1B. Activity of luciferase was measured after 1h refolding at 25 °C and shown as an average of at least three repeats ± standard deviation. (B, C) Effect of substitutions at analyzed positions on binding of IbpA from E. coli (B) and E. amylovora (C) to heat-aggregated *E*.*coli* proteins. Assay was performed as in 3A. (D-H) Effect of substitutions at analyzed positions on inhibition of Hsp70 system binding to aggregates by extant sHsps (D) Experimental scheme. (E-H) Aggregate-bound sHsps differently inhibit Hsp70 binding. BLI sensor with immobilized aggregated luciferase and aggregate bound sHsps was incubated with Hsp70 or buffer (spontaneous dissociation curve). Grey traces present Hsp70 binding to immobilized aggregates in the absence of sHsps. Results are presented as an average of at least three repeats ± standard deviation.

Above results suggest that tight sHsp binding to aggregates negatively affects subsequent Hsp70-Hsp100-dependent substrate refolding process. It is initiated by binding of the Hsp70 system (DnaK and cochaperones DnaJ and GrpE) to aggregates that requires sHsps to be outcompeted from aggregates (Zwirowski *et al*, 2017). To gain insight into the competition between sHsps and Hsp70 we modified the biolayer interferometry experiments and introduced the sensor with sHsps bound to luciferase aggregates into a buffer containing Hsp70 system (Fig. 6D). Although biolayer interferometry cannot distinguish between proteins bound to the sensor, we took advantage of the differences in the thickness of the protein layers specific for sHsp or Hsp70 binding and also in the binding kinetics. The analysis of the Hsp70 binding to the aggregates covered with sHsps clearly shows that the presence of IbpA_*E*.*amyl*_ or IbpA_*E*.*coli*_ Q66H G109D on aggregates only weakly inhibits Hsp70 binding (Fig. 6E,H). In contrast, the inhibition is much more pronounced when IbpA_*E*.*amyl*_ H67Q D110G or IbpA_*E*.*coli*_ are present on aggregates (Fig. 6F,G).

All above experiments indicate that two specific amino acids in positions 66 and 109 in ACD of IbpA proteins define the mode of IbpA activity. Glutamine 66 and glycine 109 are characteristic for IbpA proteins which bind tightly to substrates and thus are not easily outcompeted from the aggregates by Hsp70s. Such IbpAs require IbpB partner cooperation to function properly. Substitutions at these positions to histidine (position 66) and aspartic acid (position 109) allowed for the emergence of a single sHsp which binds to aggregating substrate less tightly and can be outcompeted from the aggregates by Hsp70s in the absence of IbpB.

## Discussion

In this study we traced, at the molecular level, how the two-protein sHsp system, a part of the cellular protein refolding machinery, underwent simplification in a way that its biochemical functions are performed by a single protein. In most *Enterobacterales* two sHsps (IbpA and IbpB) drive sequestration of misfolded proteins into the reactivation-prone assemblies (Obuchowski *et al*., 2019). Together, IbpA and IbpB form a functional heterodimer, in which IbpA specializes in substrate binding, preventing the substrates from creating large aggregates (sequestrase activity), while IbpB promotes sHsps dissociation from the aggregates required for subsequent Hsp70-Hsp100-dependent substrate refolding (Obuchowski *et al*., 2019; Pirog *et al*., 2021). We showed that, in parallel to the *ibpB* gene loss in *Erwiniaceae*, new functions of IbpA have emerged, i.e. a lower substrate sequestrase activity, which correlates with the efficient substrate refolding. We have identified two amino acid substitutions (Q66H and G109D) responsible for this new IbpA functionality. Selection analysis shows that these two substitutions were likely driven by positive selective pressure. This indicates that this change possibly had an adaptive character in the ancestral background. It is important to note, however, that models used for the analysis do not account for variation of synonymous substitution rate and multinucleotide substitution events, which in some cases might lead to false positive results (Lucaci *et al*, 2023). Because of that, the alternative hypothesis that substitutions Q66H and G109D occurred in the common ancestor of *Erwiniaceae* due to genetic drift, enabling the subsequent loss of *ibpB* gene, cannot be fully discounted. Functional differences observed between modern-day sHsps from *E. coli* and *E. amylovora* are at least partially defined by presence of specific amino acids in these two positions and can be diminished by their swapping between extant proteins. Their occurrence in the last common ancestor of *Erwiniaceae* IbpA resulted in a decreased affinity of ACD’s β4-β8 cleft to aggregated substrates as well as to the C–termini of the other sHsps. These interactions might be of particular importance for stabilization of sHsps on a surface of sequestered aggregated substrates leading to the formation of so-called protective shell preventing further uncontrolled aggregation. Apparent role of C-termini and ACD interaction is in agreement with earlier studies, showing that addition of free C-terminal peptide causes *E. coli* IbpA and IbpB dissociation from the outer shell of sHsp - substrate complex (Zwirowski *et al*., 2017). It was also shown that, in case of Hsp 16.6 from cyanobacterium *Synechocystis*, substitutions that slightly weakened interaction between ACD and C-termini lead to increased stimulation of luciferase refolding. However, abolishing this interaction resulted in a non-functional protein, most likely due to the loss of the sequestrase activity (Giese & Vierling, 2002). Destabilization of the protective shell of secondarily single IbpA by weakening these interactions had an effect functionally analogous to a role of IbpB in the two-protein system, facilitating IbpA dissociation from the substrate (Obuchowski *et al*., 2019). Two identified substitutions also weaken the ACD interaction with aggregated substrates which is an additional factor shifting the sHsps balance towards dissociation, a step necessary to initiate Hsp70-Hsp100 dependent disaggregation and refolding.

Our results show how ACD substitutions can fine-tune sHsp system by exerting pleiotropic effects on ACD - C-terminal peptide and ACD - substrate interactions. These relatively small changes strongly influence the effectiveness of sHsp functioning in complex process of aggregated protein rescue by molecular chaperones. It might be particularly important in the case of a conserved interaction, like the one between C - terminal peptide and ACD, when excessive changes of affinity may be detrimental to the overall protein function (Giese & Vierling, 2002). Our approach enabled us to find functional residues in sHsp system, which would not have been possible by using conventional mutagenesis and highlights the importance of using vertical approach in biochemical studies. This study closely follows the evolutionary process, in which mutations in one of the two cooperating proteins tinker it to a point it becomes independent of its partner, enabling the simplification of a more complex system by partner loss while maintaining its overall function.

Following, with molecular precision, the genetic events associated with the gene loss allowed us to answer several questions about protein family evolution. The first question concerns how the lost function is incorporated into a remaining partner protein. The results of our experiments indicate that even though the primary function of a partner protein is maintained, it is altered so that it does not interfere with the newly incorporated one. In our case substrate binding properties, as well as formation of higher-order oligomers were altered in a way to keep the sequestrase function maintained but allow for more efficient stimulation of Hsp70-Hsp100-dependent substrate refolding. The second question concerns the context-dependence of mutations in protein evolution. We successfully transplanted the mutations that appeared in *E. amylovora* into *E. coli* IbpA ortholog, artificially creating an efficient single protein system from a protein that normally needs a partner for efficient substrate refolding. In contrast to other studies (Natarajan *et al*, 2023) the context of *E*.*coli* IbpA protein did not influence the ability of IbpA_E.coli_ Q66H G109D to work without IbpB partner. This brings the question about the accessibility of adaptive solutions. In case of *E*.*coli* IbpA there was no need for adaptive changes because the gene loss did not occur in this clade, suggesting that the loss of a protein can push another one towards an adaptation, which leads to finding efficient molecular innovations.

## Materials and Methods

### Reconstruction of IbpA phylogeny

Amino acid sequences of 77 IbpA orthologs from *Enterobacterales* were obtained from NCBI and UniProt databases and aligned using Clustal Omega (Sievers *et al*, 2011). Alignment was trimmed manually. JTT+R3 was identified as the best fit model by iq-tree, using Bayesian Information Criterion and was used in the analysis (Kalyaanamoorthy *et al*, 2017; Nguyen *et al*, 2015). The phylogenetic tree was inferred using iq-tree on the basis of 328 iterations of ML search with 100 rapid bootstraps replicates (Nguyen *et al*., 2015).

### Reconstruction of ancestral IbpA amino acid sequences

Ancestral sequence reconstruction was performed on the basis of multiple sequence alignment of 77 amino acid sequences of IbpA orthologs from *Erwiniaceae* and *Enterobacreriaceae* as well as a phylogenetic tree of those orthologs (see above). Marginal reconstruction of ancestral sequences was performed with FastML program based on ML algorithm and Bayesian approach using JTT substitution matrix with gamma parameter (Ashkenazy *et al*., 2012; Cohen & Pupko, 2011; Cohen *et al*., 2008; Jones *et al*, 1992; Pupko *et al*., 2002; Simmons & Ochoterena, 2000).

Alternative ancestral sequences for AncA_0_ and AncA_1_ proteins were obtained by substituting most likely amino acid on every uncertain position (defined as a position with more than one amino acid with posterior probability ≥ 0.2) with the amino acid with the second highest posterior probability (Eick *et al*., 2017).

### Analysis of natural selection

Analysis of natural selection was performed using codeml. First, Pal2Nal was used to obtain codon alignment based on the multiple sequence alignment of amino acid sequences of IbpA orthologs from *Enterobacterales* (see above) as well as corresponding nucleotide sequences obtained from NCBI database. Resulting codon alignment was then trimmed manually and used together with the phylogenetic tree obtained earlier (see above) for the selection analysis.

For branch model analysis, models M0 (null hypothesis) and Two – ratio (with either AncA_0_ -AncA_1_ branch or *Erwiniaceae* clade as foreground) were used. For branch – site model analysis, models A null (null hypothesis) and A were used, with foreground branches selected as above. Statistical significance of different models was estimated with Likelihood Ratio Test (LRT) (Jeffares *et al*., 2015; Yang, 1998, 2007; Yang & Nielsen, 2002).

### Protein Purification

#### Purification of IbpA proteins

pET3a plasmids containing *ancA*_*0*_, *ancA*_*1*_, *ancA*_*0*_ *+7, ancA*_*0 alt_all*_, *ancA*_*1alt_all*_ and *ibpA*_*Ea*_ genes were ordered from GeneScript. Point mutations were introduced using site - directed mutagenesis and confirmed by sequencing. Proteins were overproduced in *E. coli* BL21(DE3). Cells were then lysed by sonication in Qsonica sonicator (13% amplitude, 2 min 30 s process time, 15 s pulse-ON time, 45 s pulse-OFF time) in lysis buffer L1 (50 mM Tris pH 7.5, 50 mM NaCl, 5 mM EDTA, 10 % glycerol, 5mM β-mercaptoethanol). Insoluble fraction containing proteins of interest was separated by centrifugation (75 000 x g, 30 min, 4°C) and resolubilized in buffer A (40 mM Tris pH 7.5, 50 mM NaCl, 10% glycerol, 5 mM β-mercaptoethanol, 6M urea) and then centrifuged (75 000 x g, 30 min, 4°C). Supernatant was loaded on Q – Sepharose chromatography column equilibrated with buffer A and eluted in 50 mM - 500 mM NaCl gradient. Fractions containing proteins of interest were then dialyzed to buffer B (40 mM Tris pH 8.5, 50mM NaCl, 10% glycerol, 5mM β-mercaptoethanol) and loaded on Q – Sepharose chromatography column equilibrated with buffer B. Flow-through fraction was collected and dialyzed to buffer C (50 mM Tris pH 7.5, 150 mM KCl, 5% (v/v) glycerol, 5 mM β-mercaptoethanol).

#### Purification of ACD domains

ACDs of IbpA_*Ec*_,, AncA_0_ and AncA_0_ Q66H G109D were purified as described previously (Pirog *et al*., 2021) and as a final step dialyzed to buffer G (50 mM Tris pH 7.5, 150mM KCl, 5mM β-mercaptoethanol).

#### Purification of His_6_ – SUMO and His_6_ –SUMO-C-terminal peptide of AncA_0_ construct

His_6_ - SUMO was purified using the Champion™ pET SUMO Expression System. pET28a plasmid containing gene encoding His_6_ –SUMO fused with C-terminal peptide of AncA_0_ (PEAMKPPRIEIN) was ordered from GeneScript. Proteins were overproduced in *E. coli* BL21(DE3). Cells were then lysed by sonication in Qsonica sonicator (20% amplitude, 2 min process time, 5s pulse-ON time, 10s pulse-OFF time) in lysis buffer L2 (40 mM Tris pH 7.5, 100 mM NaCl, 10 % glycerol, 10 mM imidazole, 2 mM β-mercaptoethanol). Insoluble fractions were separated by centrifugation for 30 min at 70 000 x g and supernatants, containing proteins of interest, were incubated for 1h with Ni-NTA resin equilibrated with buffer L2. Resins were then washed with the buffer D (40 mM Tris pH 7.5, 100 mM NaCl, 10 % glycerol, 40 mM imidazole, 2mM β-mercaptoethanol) Proteins of interest were eluted from the columns with the buffer E (40 mM Tris pH 7.5, 100 mM NaCl, 10 % glycerol, 400 mM imidazole, 2mM β-mercaptoethanol) and then dialyzed to buffer C (as above).

DnaK, DnaJ, GrpE, ClpB and IbpA_*Ec*_ proteins were purified as described previously (Ratajczak *et al*., 2009). IbpB_*Ec*_ protein was purified as described previously (Pirog *et al*., 2021). His-tagged luciferase used for BLI measurements was purified as described previously (Obuchowski *et al*., 2019).

Purity of purified proteins was assessed with SDS-PAGE electrophoresis with Coomassie Blue staining. Protein concentrations were measured using Bradford reaction, with Bovine Serum Albumin as a standard. In the case of His_6_ – SUMO fused with C-terminal peptide of AncA_0_, concentration was measured with SDS-PAGE electrophoresis with Coomassie Blue staining coupled with densitometric analysis with Bovine Serum Albumin used as a standard.

OuantiLum® Recombinant Luciferase was purchased from Promega. Creatin Kinase from rabbit muscle was purchased from Sigma Aldrich.

### Luciferase refolding assay

1,5 μM recombinant firefly luciferase in buffer F (50 mM Tris pH 7.5, 150 mM KCl, 20 mM MgCl_2_, 2.5 mM DTT) was denatured by incubation for 10 min at 44°C alone or in the presence of 10 μM sHsps (3uM IbpA_*Ec*_ *+* 7μM IbpB_*Ec*_ in the case of two-protein system from *E. coli*). Denatured luciferase was then incubated at 25 °C with Hsp70 system (1 μM DnaK, 0.4 μM DnaJ and 0.3 μM GrpE), 2 μM ClpB, and ATP regeneration system (5 mM ATP, 0.1 mg/ml creatine kinase and 18 mM creatine phosphate). For experiment presented in Fig.3 – figure supplement 3D, higher concentration of the Hsp70 system was used (2 μM DnaK, 0.8 μM DnaJ and 0.6 μM GrpE). At different timepoints luciferase activity was measured with GLOMAX™ 20/20 luminometer, using the Luciferase Assay System from Promega. Results are presented as averages of at least three independent repeats ± standard deviation.

### DLS measurements

Dynamic Light Scattering measurements were performed using Malvern Instruments ZetaSizer Nano S instrument, at 40 μl sample volume, scattering angle 173° and wavelength of 633 nm. For every measurement, minimum ten subsequent series of ten 10-s runs were averaged and particle size distribution was calculated by fitting to 70 size bins between 0.4 and 10,000 nm, as previously described (Zwirowski *et al*., 2017).

For reversible deoligomerization assay, size of oligomers formed by 10 μM sHsps in buffer F (50 mM Tris pH7.5, 150 mM KCl, 20 mM MgCl_2_, 2.5 mM DTT) were measured by DLS. First measurement was performed at 25°C and then the sample was heated to 44°C, cooled to 25°C, heated to 44°C and cooled to 25°C, with measurements performed after each change in temperature. Results are presented as size distribution by volume.

For measuring influence of substitutions on oligomer formation, size of oligomers formed by either 10 μM AncA_0_ or 10 μM AncA_0_ Q66H G109D in buffer F (as above) was measured by DLS at 25, 27, 29, 31, 33, 35, 37, 39, 41, 43 and 45°C. Results were presented as size distribution by intensity (for temperatures 25°C, 35°C and 45°C) or as an average hydrodynamic diameter corresponding to maximum of a dominant peak of size distribution by volume plotted against temperature ± standard deviation.

For assembly formation (sequestrase) assay, 1.5 μM firefly luciferase in buffer F was denatured alone or in the presence of different sHsp concentrations by incubation at 44°C for 10 min. Size of obtained luciferase aggregates was then measured by DLS at 25°C. Results are presented as an average hydrodynamic diameter of measured particles weighted by intensity (Z-average) ± standard deviation.

### Biolayer interferometry (BLI) measurements

sHsps interactions with aggregated luciferase or aggregated *E. coli* lysate were measured using Octet ® K2 system. Anchoring layer of his-tagged luciferase was attached to Octet® NTA Biosensors by 5 min incubation in 0.6 mg/ml his-tagged luciferase in denaturing conditions in buffer UF (50 mM Tris pH 7.5, 4.5 M urea, 150 mM KCl, 20 mM MgCl_2_, 2.5 mM DTT) at 25 °C with 350 rpm shaking. Sensors were then incubated for 5 min as above in buffer H (50 mM Tris pH 7.5, 150 mM KCl, 20 mM MgCl_2_, 5 mM β-mercaptoethanol) to remove urea and unbound luciferase. The protein aggregate was then formed on the sensor by incubation for 10 min in 0,5 mg/ml his-tagged luciferase or 0.2 mg/ml *E. coli* lysate in buffer H at 44°C (in case of the luciferase) or 55 °C (in case of the lysate). Sensors were then again incubated in buffer H for 5 min at 25 °C with 350 rpm shaking to remove excess protein. Sensors with attached aggregate were placed in the Octet® system in H buffer for 60 s baseline measurement and then placed for 1h in 5 μM sHsp solution in H buffer to measure sHsps association. Sensors were then moved for 1h to buffer H to measure protein dissociation. Measurements were performed with 1000 rpm shaking at 44°C (in case of full - length sHsps) or at 25°C (in case of ACDs). Full-length proteins were preincubated at 44°C for 10 min before measurement.

sHsps displacement by Hsp70 system was measured using ForteBio® BLItz. Sensors with attached aggregates were prepared as described above, with buffer F (50 mM Tris pH7.5, 150 mM KCl, 20 mM MgCl_2_, 2.5 mM DTT) instead of buffer H. Baseline biolayer was measured for 60 s. Sensors were then placed in 5 μM sHsp solution in buffer F, previously preincubated for 10 min at 44°C. sHsps association was measured for 10 min. Sensors were then moved to either buffer F or Hsp70 system in buffer F (0.7 μM DnaK, 0.28 μM DnaJ, 0.21 μM GrpE, 5 mM ATP, 0.1 mg/ml creatine kinase, 18 mM creatine phosphate). Hsp70 system binding and sHsps dissociation were measured for 1h. Measurements were performed at room temperature with 2000 rpm shaking.

ACD interactions with C –terminal peptide were measured using Octet® K2 system. Octet® NTA Biosensors were placed in buffer C (50mM Tris pH 7.5, 150 mM KCl, 5% glycerol, 5 mM β-mercaptoethanol) and baseline signal was measured for 60 s. Sensors were then placed in 2.5 μM His_6_ -SUMO-C-peptide solution in buffer C and incubated for 15 min. Surplus His_6_ -SUMO-C-peptide was then removed by incubation in G buffer for 15 min. Sensors were then moved to ACD solution and association was measured for 48 min. After that, ACD dissociation was measured in buffer G for 10 min. Measurements were performed with 1000 rpm shaking at 25°C. Measurements were performed for different ACD concentrations in triplicates. Biolayer thickness at the end of association stage was corrected for the nonspecific binding using control substituting His_6_ -SUMO for His_6_ -SUMO-C-peptide and converted to fraction bound by min-max scaling between 0 and 1 using minimal and maximal triplicate averages of biolayer thickness. To determine dissociation constant fraction bound as a function of ACD concentration was fitted to the Hill equation using SciPy implementation (Virtanen *et al*, 2020) of dogbox algorithm:

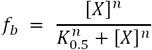

Where [X] is the total ACD concentration, K_0.5_ – ACD concentration required to reach half-maximum binding at equilibrium, n – Hill coefficient, f_b_ - fraction bound. Standard deviations of fitted K_0.5_ and n values were derived from the diagonal of optimized parameters covariance matrix.

### Molecular Dynamics (MD)

All MD simulations were performed using Gromacs 2019.2 (Van Der Spoel *et al*, 2005) and CHARMM36-jul2021 as a force field (Huang *et al*, 2017). Simulations were performed in the isothermal-isobaric (NPT) ensemble, where temperature was kept at 310 K using v-rescale thermostat (Bussi *et al*, 2007) with a time constant of 0.1 ps and the pressure was held at 1 bar using Parinello-Rahman barostat (Parrinello & Rahman, 1981). Lennard-Jones potential with a cut-off of 1.0 nm was used to describe Van der Waals interactions. Computation of long-range electrostatic interactions was performed using the particle mesh Ewald (PME) method (Essmann *et al*, 1995) with a Fourier grid spacing of 0.12 nm and a real space cutoff of 1.0 nm. Bonds between hydrogen and protein heavy atoms were constrained by P-LINCS (Hess *et al*, 1997) and water molecules geometry was constrained by SETTLE (Miyamoto & Kollman, 1992). Integration of equations of motion was performed by leap-frog algorithm (Van Gunsteren & Berendsen, 1988) with a time step of 2 fs. Periodic boundary conditions were applied in all dimensions.

The initial conformation of AncA_0_ ACD - C-terminal peptide complex was predicted by ColabFold implementation (Mirdita *et al*, 2022) of AlphaFold-Multimer (Richard *et al*, 2022) Substitutions Q66H and G109D were introduced to the complex using PyMol Mutagenesis Wizard (http://www.pymol.org). Complexes were then placed in rhombic dodecahedral boxes measuring 10.4 nm in all dimensions and solvated by CHARMM-modified TIP3P water model (Jorgensen *et al*, 1983). Concentrations of sodium and chloride ions were adjusted to 0.15 M and net zero charge of the system. Each system was subjected to 3-step energy minimization protocol, where during the first step protein conformation was constrained, during the second step constraint was reduced to protein backbone only and during the third step positions of all protein heavy atoms were restrained with a force constant of 1000 kJ*mol^-1^*nm^-1^. Minimized systems were equilibrated for 10 ns while positions of protein backbone atoms were kept constrained, equilibration was then continued without constraints for further 500 ns. The first 100 ns of equilibration was discarded, and the rest was used for ACD – C-terminal peptide contact determination using GetContacts tool (https://getcontacts.github.io/). Contact between interfacial residues was defined as any of the following types of interaction: hydrogen bond, ionic, π-stacking, π-cation or van der Waals between purely hydrophobic residues. The default GetContacts interaction criteria were used for all interaction types except for hydrogen bond detection, where a more stringent 30° cutoff for hydrogen-donor-acceptor angle was used. Contact heatmaps were prepared using seaborn (Waskom, 2021).

The representative conformations of ACD – C-terminal peptide complexes were chosen by clustering the last 400 ns of equilibrium MD trajectories (frames spaced every 0.5 ns) using “gmx cluster” tool and Jarvis-Patrick clustering method (Jarvis & Patrick, 1973). RMSD cutoff of 0.2 nm, was used for Jarvis-Patrick algorithm and conformations possessing at least 3 neighbors in common out of 15 closest conformations were assigned to the same cluster. RMSD calculation was based on coordinates of heavy backbone atoms of the C-terminal peptide interacting ACD monomer without dimerization loop (residues 40-74 and 95 to 126) and stably interacting region of the C-terminal peptide (residues 132-137). In the case of both simulated complexes the biggest cluster contained more than 90% of simulation frames (92.3% for AncA0 ACD complex and 93.3% for Anca0 Q66H G109D complex) and its middle frame was chosen as representative conformation. Visual Molecular Dynamics (VMD) (William *et al*, 1996) and Blender (https://www.blender.org/) were used for structure visualization.

## Supporting information

supplementary file 1 - Multiple Sequence Alignment of Enterobacterales IbpA orthologs

supplementary file 2 - phylogenetic tree of Enterobacterales IbpA orthologs in Newick format

supplementary file 3 - posterior probability statistics

supplementary file 4 - branch and branch - site model statistics

## Acknowledgements

This work was supported by a grant of the Polish National Science Centre (OPUS 17 2019/33/B/NZ1/00352). We gratefully acknowledge Poland’s high-performance Infrastructure PLGrid (HPC Centers: ACK Cyfronet AGH, PCSS, CI TASK, WCSS) for providing computer facilities and support. We thank Prof. Max Telford, Prof. Jaroslaw Marszalek and Dr. Agnieszka Kłosowska for helpful discussions.

## Additional Files

### Figure supplements

**Supplementary file 1** Multiple sequence alignment of *Enterobacterales* IbpA orthologs used for phylogenetic analysis and ancestral reconstruction in fasta format

**Supplementary file 2 Phylogenetic tree of *Enterobacterales* IbpA protein family in newick format** (see figure 2)

**Supplementary file 3 Posterior probability statistics for ancestral sequence reconstruction of AncA**_**0**_ **and AncA**_**1**_ **nodes:** For each position amino acids reconstructed with posterior probability higher than 0.2 are shown. Single – letter symbols of reconstructed amino acids are followed by posterior probability of reconstruction (in brackets). Positions at which most likely amino acid differ between AncA_0_ and AncA_1_ are marked in bold and italics. Posterior probabilities estimated using FastML program based on Maximum Likelihood and Empirical Bayes method.

**Supplementary file 4 Statistics for selection analysis:** A) Branch model statistics for IbpA orthologs from *Enterobacterales*. Models assumed either branch between nodes AncA_0_ and AncA_1_ or entire *Erwiniaceae* clade as foreground; NS – Not significant. B) Branch - site model statistics for IbpA orthologs from *Enterobacterales*. Models assumed either branch between nodes AncA_0_ and AncA_1_ or entire *Erwiniaceae* clade as foreground; NS – Not significant

