## Supplementary material for "Evolution towards simplicity in bacterial small heat shock protein system": Figure_supplements

**Figure3 – figure supplement 1**

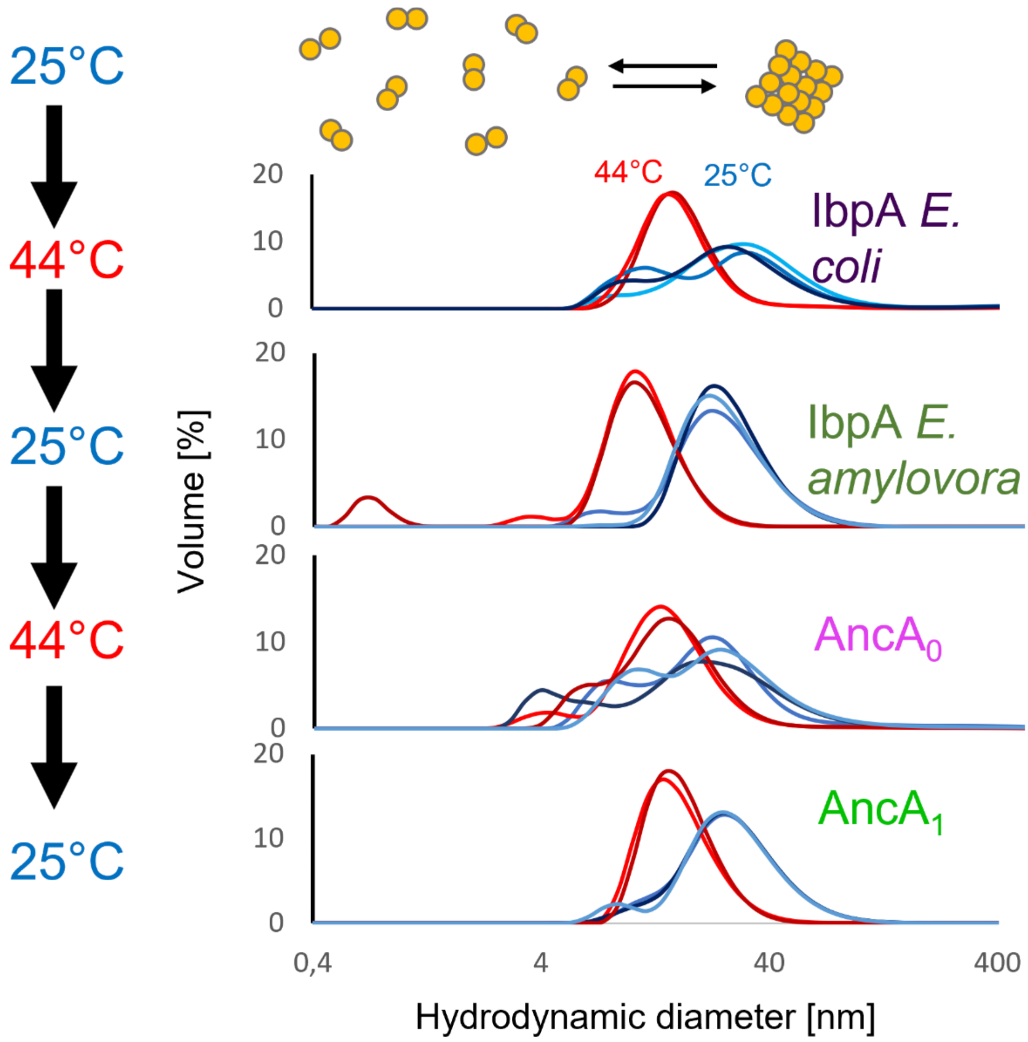


**Fig. 3–figure supplement 1. Both reconstructed proteins reversibly deoligomerize at heat shock temperature, similarly to extant proteins:** Hydrodynamic diameter distributions of sHsp oligomers incubated subsequently at 25°C (blue graphs) and 44°C (red graphs). Size distributions were measured by DLS.

**Figure3 – figure supplement 2**

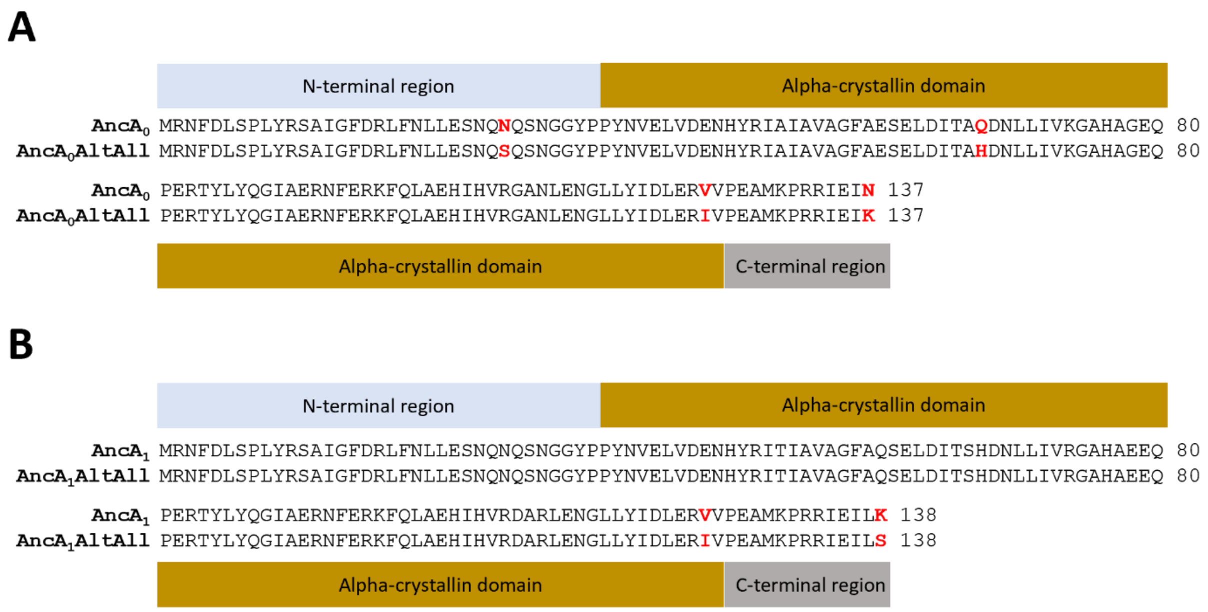


**Figure 3 – Figure supplement 2. Amino acid sequences of ML and AltAll variants of reconstructed ancestral proteins:** $AncA_0$ , $AncA_1$ – ML variants; $AncA_0$AltAll, $AncA_1$AltAll – AltAll variants. Positions differing between ML and AltAll variants are marked in red. (A) Alignment of $AncA_0$ and $AncA_0$AltAll sequences. (B) Alignment of $AncA_1$ and $AncA_1$AltAll sequences.

Figure3 – figure supplement 3

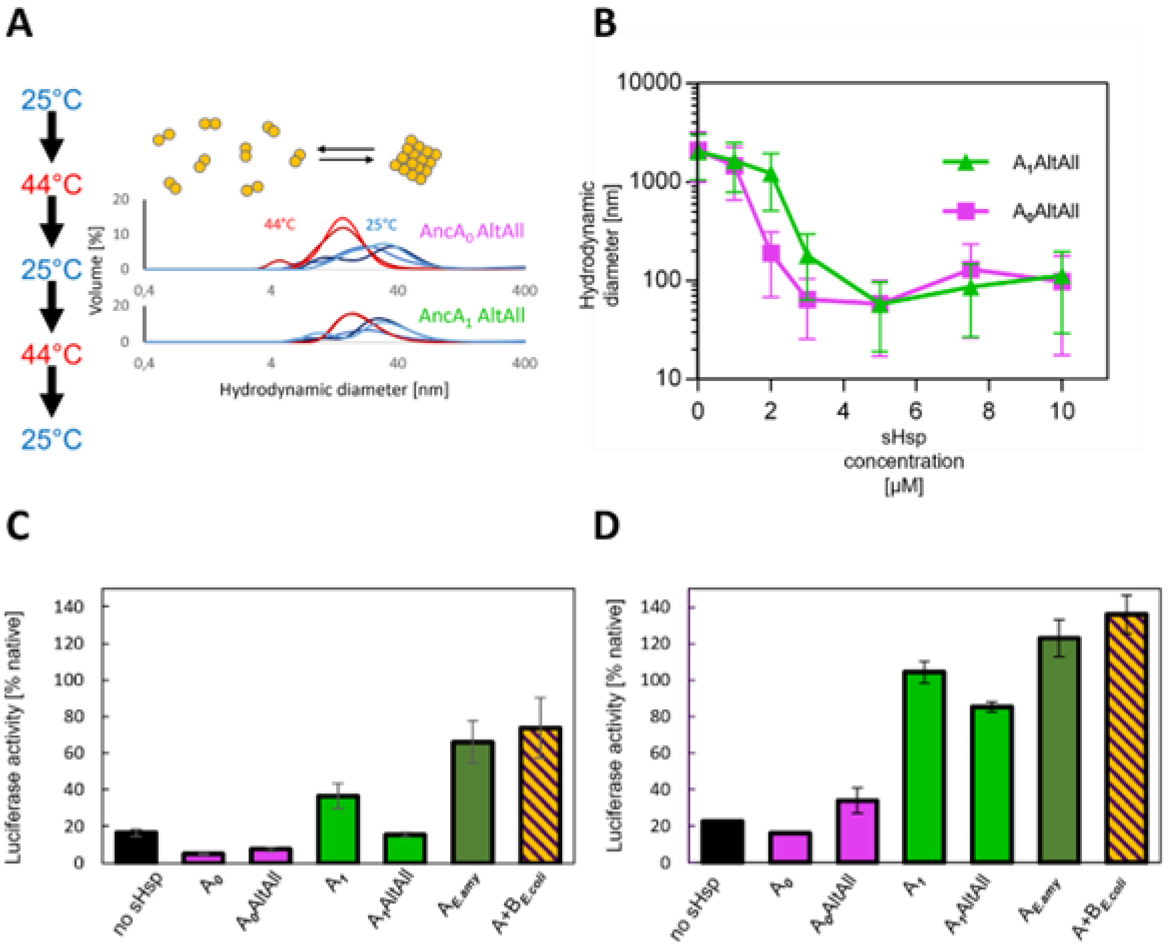


**Fig.3 – figure supplement 3. AltAll variants of AncA$_0$ and AncA$_1$ have similar properties to ML variants:** (A) Reversible deoligomerization at heat shock temperature; measurement were performed as in Fig. 3 – figure supplement 1. (B) Sequestrase activity. Luciferase sequestration assay was performed as in Fig. 3B. Results are shown as an average hydrodynamic diameter ± standard deviation. (C-D) Ability to stimulate Hsp70-Hsp100 - dependent luciferase disaggregation; Luciferase refolding assay was performed as in Fig. 1. Activity of luciferase was measured after 1h refolding at 25 °C and shown as an average of at least three repeats ± standard deviation; assays were performed at two concentrations of Hsp70 system: (C) standard (1 μM DnaK, 0.4 μM DnaJ 0.3 μM GrpE) or (D) increased (2 μM DnaK, 0.8 μM DnaJ 0.6 μM GrpE)

### Figure 4 – figure supplement 1

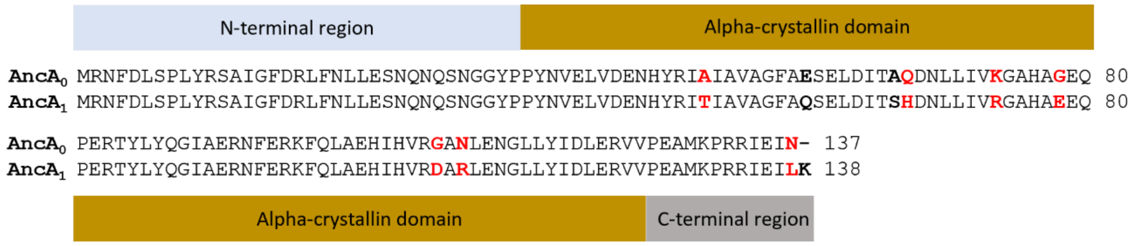


**Fig. 4 – figure supplement 1 Amino acid sequence differences between AncA$_0$ and AncA$_1$:** Differing positions used in further analysis were marked in red, differing positions omitted from further analysis due to low conservation in extant *Erwiniaceae* were marked in bold.

### Figure 4 – Figure supplement 2

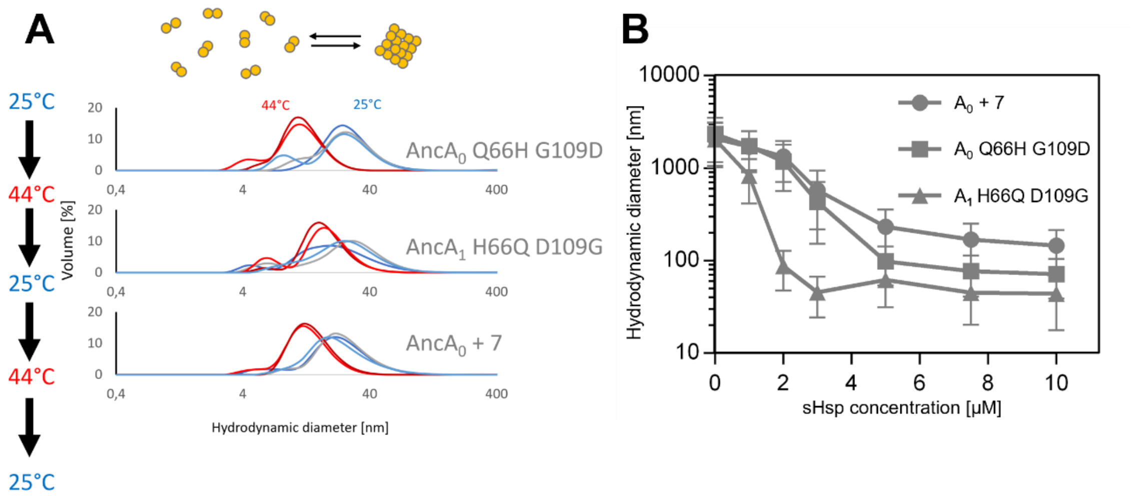


**Fig. 4 – figure supplement 2. AncA$_0$ Q66H G109D, AncA$_0$+7 and AncA$_1$ H66Q D109G exhibit sequestrase activity and reversibly deoligomerize at heat shock temperature:** (A) Reversible deoligomerization at heat shock temperature; measurements were performed as in Fig. 3–figure supplement 2. (B) Sequestrase activity; luciferase sequestration assay was performed as in Fig. 3B; results are shown as an average hydrodynamic diameter ± standard deviation.

### Figure 5 – figure supplement 1

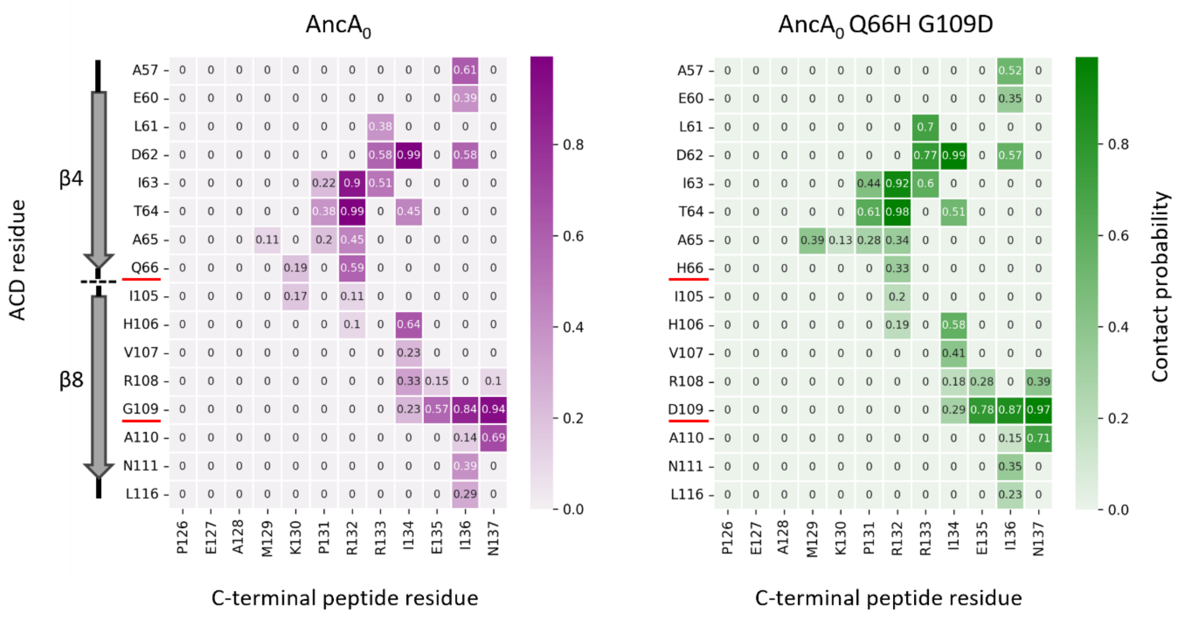


**Fig. 5 – figure supplement 1. AncA$_0$ residues 66 and 109 contact the C-terminal peptide, although overall contact pattern remains similar upon introduction of Q66H G109D substitutions:** Heatmaps illustrating contact probabilities, derived from last 400 ns of equilibrium MD simulations, between residues of the ACD (vertical-axis) and the C-terminal peptide (horizontal axis) for AncA0 (left) and AncA0 Q66H G109D (right). Number within each cell of the heatmap and cell shading represent contact probability. Schematic next to the vertical axis of AncA0 heatmap represents positioning of the ACD residues within secondary structure elements. Substituted ACD residues are underlined.

**Figure 5 – figure supplement 2**

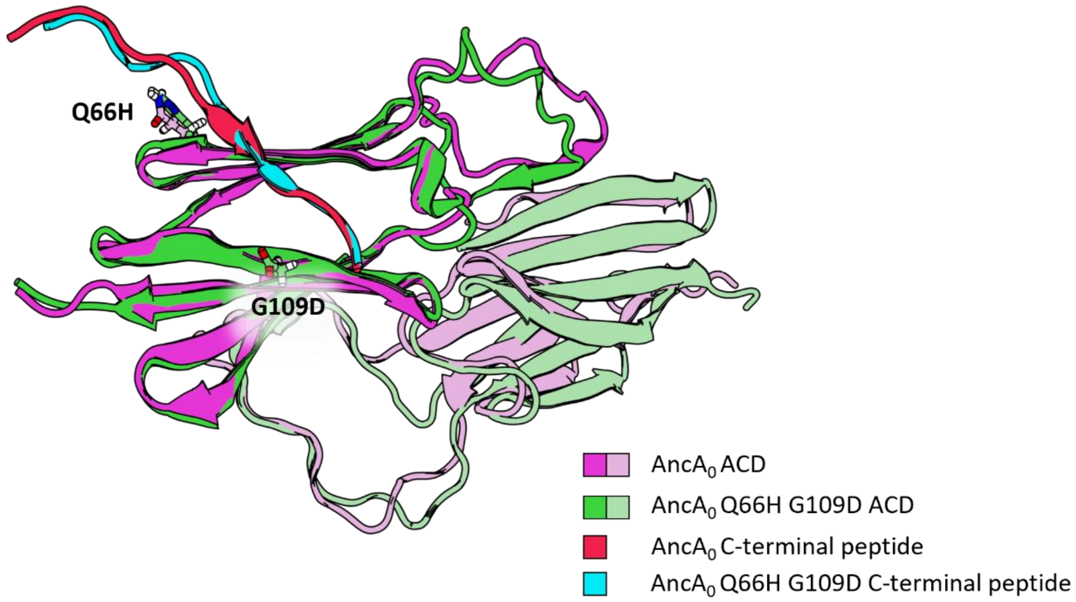


**Fig. 5 – figure supplement 2 Comparison of the structural models of AncA0 (purple) and AncA0 Q66H G109D (green) ACD dimers complexed with the C-terminal peptides:** Models represent middle structures of largest clusters obtained from equilibrium molecular dynamics simulation trajectories based on the AlphaFold2 prediction. Models were superimposed on the backbone heavy atoms of the C-terminal-peptide interacting ACD monomer without dimerization loop (residues 40-74 and 95 to 126) and the stably interacting C-terminal peptide fragment (residues 132-137), RMSD of the superimposed regions backbone atoms – 0.81 Å, RMSD of the whole complex backbone atoms – 3.42 Å. Substituted residues in positions 66 and 109 are shown in licorice representation

**Figure 5 – figure supplement 3**

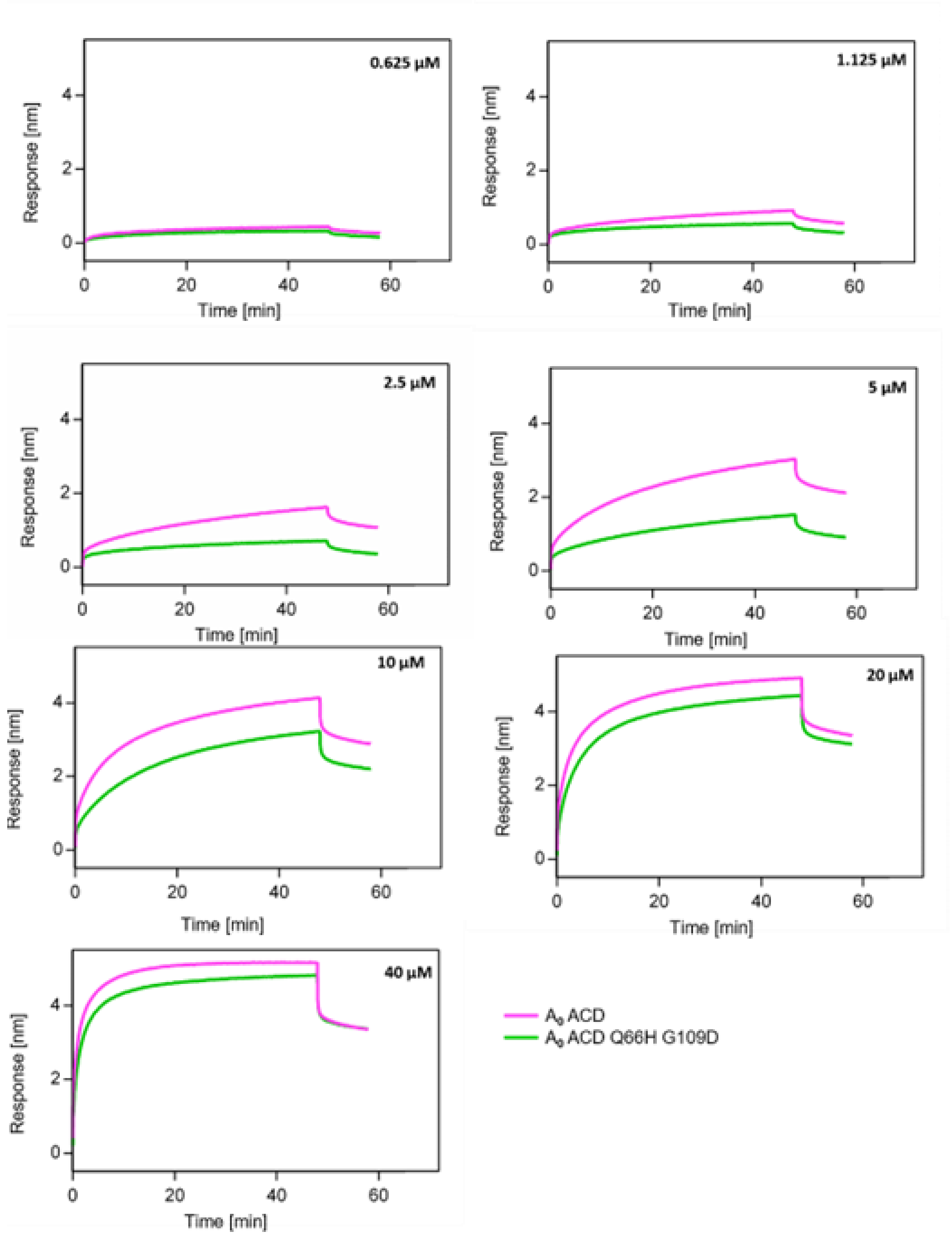


**Fig. 5 – figure supplement 3. Substitutions Q66H G109D decrease affinity of AncA$_0$ ACD to AncA$_0$ C – terminal peptide:** Representative binding curves for different concentrations of AncA$_0$ ACD and AncA$_0$ Q66H G109D ACD interacting with AncA$_0$ C- terminal peptide; background was not subtracted. Peptide was immobilized on BLI NTA sensor via His$_6$-Sumo tag.

**Figure 5 – figure supplement 4**

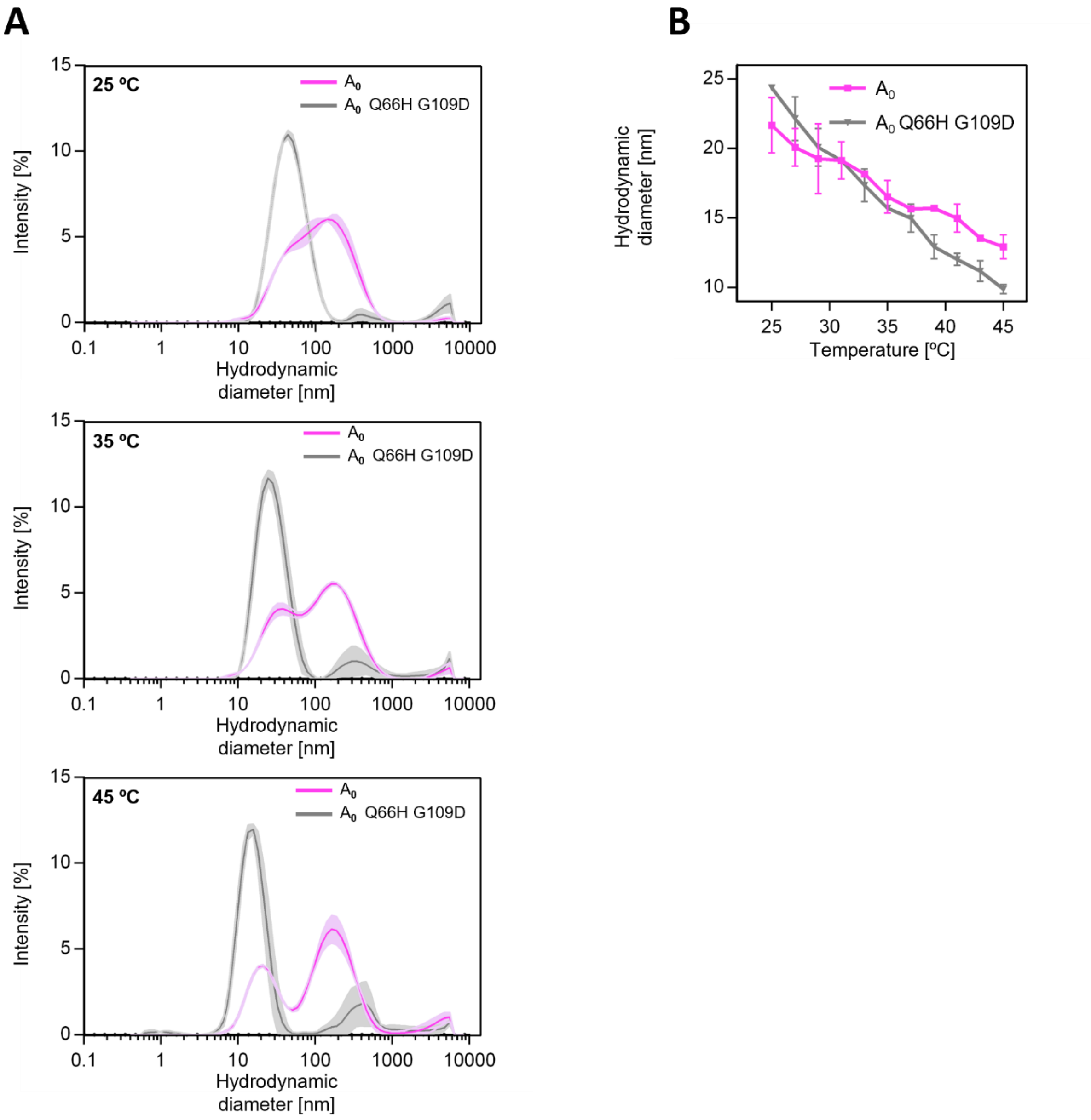


**Fig. 5–figure supplement 4. Substitutions Q66H G109D influence AncA$_0$ oligomerization:** (A) Hydrodynamic diameter distributions shown as DLS signal intensity for better visualization purposes of AncA$_0$ and AncA$_0$ Q66H G109D measured at 25, 35 and 45 °C. Size distributions were measured by DLS. Results are shown as an average of three repeats ± standard deviation. (B) Changes of hydrodynamic diameter representing maximum of the dominant peak of hydrodynamic diameter distribution by volume with temperature. Results are shown as an average of three repeats ± standard deviation.

**Figure 5 – figure supplement 5**

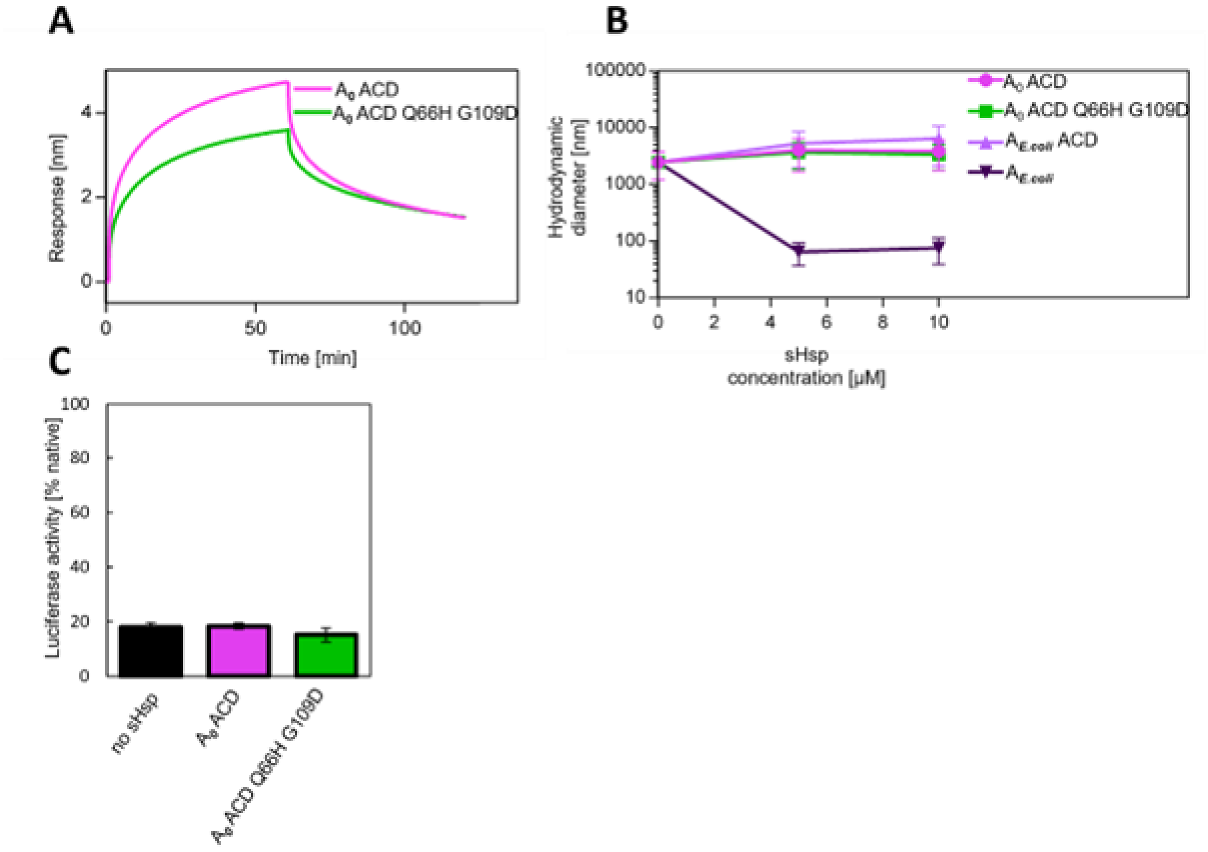


**Fig. 5 – figure supplement 5. AncA$_0$ ACD and AncA$_0$ ACD Q66H G109D can bind to aggregated luciferase, but does not exhibit sequestrase activity or ability to stimulate luciferase refolding:** (A) Effect of Q66H G109D substitutions on AncA$_0$ ACD affinity to aggregated luciferase; analysis was performed as in Fig. 3A. (B) Ability of AncA$_0$ ACD and AncA$_0$ Q66H G109D ACD to sequester aggregating luciferase. Results were compared to ACD of extant *E. coli* IbpA, as well as full – length extant *E. coli* IbpA. Analysis was performed as in Fig. 3B. Results are shown as an average hydrodynamic diameter ± standard deviation. (C) Ability of AncA$_0$ ACD and AncA$_0$ Q66H G109D ACD to stimulate Hsp70-Hsp100 - dependent luciferase disaggregation. Luciferase refolding assay was performed as in Fig. 1. Activity of luciferase was measured after 1h refolding at 25 °C and shown as an average of at least three repeats ± standard deviation.

**Figure 6 – figure supplement 1**

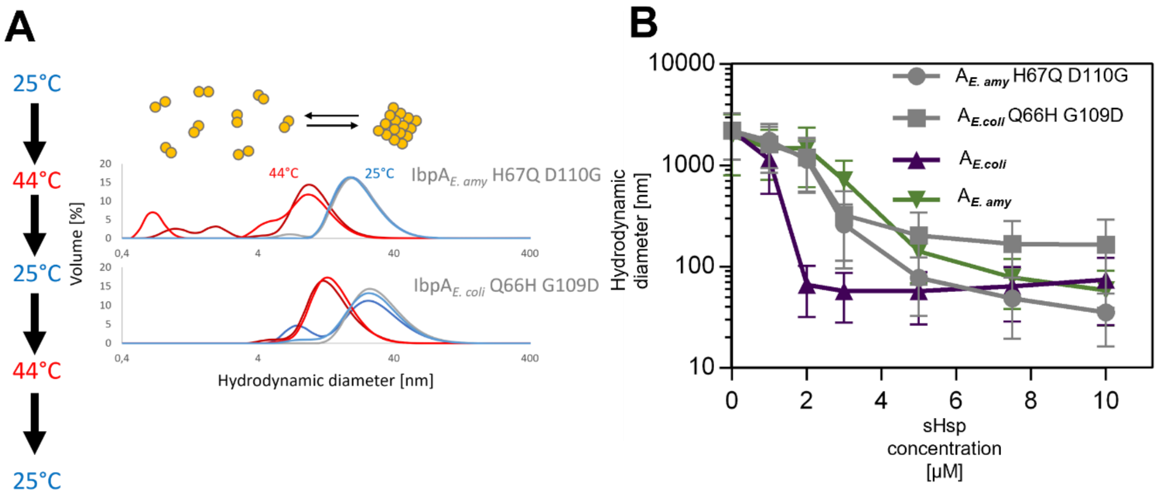


**Fig. 6–figure supplement 1. IbpA$_{E.coli}$ H66Q G109D and IbpA$_{E.amyl}$ H67Q D110G exhibit sequestrase activity and reversibly deoligomerize at heat – shock temperature:** (A) Reversible deoligomerization of IbpA$_{E.coli}$ H66Q G109D and IbpA$_{E.amyl}$ H67Q D110 at heat shock temperature. Measurements were performed as in Fig. 3 – figure supplement 2. (B) Sequestrase activity of IbpA$_{E.coli}$ H66Q G109D and IbpA$_{E.amyl}$ H67Q D110 at heat shock temperature. Measurements were performed as in Fig. 3B. Results are shown as an average hydrodynamic diameter ± standard deviation.
