## supplementary file 3 - posterior probability statistics for "Evolution towards simplicity in bacterial small heat shock protein system"

**Supplementary file 3 Posterior probability statistics for ancestral sequence reconstruction of $AncA_0$ and $AncA_1$ nodes:** For each position amino acids reconstructed with posterior probability higher than 0.2 are shown. Single – letter symbols of reconstructed amino acids are followed by posterior probability of reconstruction (in brackets). Positions at which most likely amino acid differ between $AncA_0$ and $AncA_1$ are marked in bold and italics. Posterior probabilities estimated using FastML program based on Maximum Likelihood and Empirical Bayes method.

| Position Number | Posterior probabilities | |
|---|---|---|
| | **$AncA_0$** | **$AncA_1$** |
| **1** | M (0.999926) | M (0.999866) |
| **2** | R (0.999613) | R (0.999411) |
| **3** | N (0.999721) | N (0.999542) |
| **4** | F (0.999497) | F (0.999873) |
| **5** | D (0.999313) | D (0.999001) |
| **6** | L (0.999064) | L (0.999692) |
| **7** | S (0.998966) | S (0.997905) |
| **8** | P (0.99996) | P (0.999935) |
| **9** | L (0.999823) | L (0.999728) |
| **10** | Y (0.999937) | Y (0.999902) |
| **11** | R (0.999613) | R (0.999411) |
| **12** | S (0.999127) | S (0.998633) |
| **13** | A (0.999325) | A (0.99838) |
| **14** | I (0.998482) | I (0.997912) |
| **15** | G (0.999955) | G (0.999926) |
| **16** | F (0.999931) | F (0.99989) |
| **17** | D (0.999313) | D (0.999001) |
| **18** | R (0.999613) | R (0.999411) |
| **19** | L (0.999823) | L (0.999728) |
| **20** | F (0.999754) | F (0.97628) |
| **21** | N (0.999721) | N (0.999542) |
| **22** | L (0.999812) | L (0.999727) |
| **23** | L (0.9998) | L (0.999727) |
| **24** | E (0.999305) | E (0.999025) |
| **25** | S (0.961991) | S (0.995727) |
| **26** | N (0.988107) | N (0.999069) |
| **27** | Q (0.999869) | Q (0.999782) |
| **28** | N (0.620376) S (0.354683) | N (0.671642) |
| **29** | Q (0.999869) | Q (0.999779) |
| **30** | S (0.999045) | S (0.998513) |
| **31** | N (0.999721) | N (0.999542) |
| **32** | G (0.999955) | G (0.999926) |
| **33** | G (0.999955) | G (0.999926) |
| **34** | Y (0.999937) | Y (0.999902) |
| **35** | P (0.99996) | P (0.999935) |
| **36** | P (0.99996) | P (0.999935) |

| | | |
|---|---|---|
| **37** | Y (0.999937) | Y (0.999902) |
| **38** | N (0.999721) | N (0.999542) |
| **39** | V (0.998095) | V (0.997517) |
| **40** | E (0.999305) | E (0.999025) |
| **41** | L (0.999823) | L (0.999728) |
| **42** | V (0.998095) | V (0.997517) |
| **43** | D (0.999279) | D (0.998143) |
| **44** | E (0.999129) | E (0.999009) |
| **45** | N (0.999721) | N (0.999542) |
| **46** | H (0.999623) | H (0.999857) |
| **47** | Y (0.999937) | Y (0.999902) |
| **48** | R (0.999613) | R (0.999411) |
| **49** | I (0.998482) | I (0.997912) |
| **50** | ***A (0.986377)*** | ***T (0.983415)*** |
| **51** | I (0.998482) | I (0.997911) |
| **52** | A (0.999353) | A (0.999) |
| **53** | V (0.998095) | V (0.997517) |
| **54** | A (0.999353) | A (0.999) |
| **55** | G (0.999955) | G (0.999926) |
| **56** | F (0.999931) | F (0.99989) |
| **57** | A (0.999353) | A (0.998999) |
| **58** | ***E (0.993163)*** | ***Q (0.991222)*** |
| **59** | S (0.980818) | S (0.993494) |
| **60** | E (0.999305) | E (0.999025) |
| **61** | L (0.999823) | L (0.999728) |
| **62** | D (0.94017) | D (0.991033) |
| **63** | I (0.998482) | I (0.997912) |
| **64** | T (0.999463) | T (0.999145) |
| **65** | ***A (0.976453)*** | ***S (0.978346)*** |
| **66** | ***Q (0.703474) H (0.296138)*** | ***H (0.994492)*** |
| **67** | D (0.999258) | D (0.998997) |
| **68** | N (0.999721) | N (0.999542) |
| **69** | L (0.999261) | L (0.981563) |
| **70** | L (0.999823) | L (0.999728) |
| **71** | I (0.997518) | I (0.997483) |
| **72** | V (0.998094) | V (0.997516) |
| **73** | ***K (0.940376)*** | ***R (0.984241)*** |
| **74** | G (0.999955) | G (0.999926) |
| **75** | A (0.998624) | A (0.998938) |
| **76** | H (0.999878) | H (0.999865) |
| **77** | A (0.997391) | A (0.889298) |
| **78** | ***G (0.977685)*** | ***E (0.991155)*** |
| **79** | E (0.999217) | E (0.999017) |
| **80** | Q (0.99967) | Q (0.999775) |
| **81** | P (0.752893) | P (0.8025) |

| | | |
|---|---|---|
| **82** | E (0.998753) | E (0.998985) |
| **83** | R (0.999607) | R (0.999292) |
| **84** | T (0.987703) | T (0.804431) |
| **85** | Y (0.999937) | Y (0.999902) |
| **86** | L (0.999823) | L (0.999728) |
| **87** | Y (0.999937) | Y (0.999902) |
| **88** | Q (0.999869) | Q (0.999782) |
| **89** | G (0.999955) | G (0.999926) |
| **90** | I (0.998482) | I (0.997912) |
| **91** | A (0.999353) | A (0.999) |
| **92** | E (0.999305) | E (0.999025) |
| **93** | R (0.999613) | R (0.999411) |
| **94** | N (0.999721) | N (0.999542) |
| **95** | F (0.999931) | F (0.99989) |
| **96** | E (0.999305) | E (0.999025) |
| **97** | R (0.999613) | R (0.999411) |
| **98** | K (0.999614) | K (0.99943) |
| **99** | F (0.999931) | F (0.99989) |
| **100** | Q (0.999869) | Q (0.999782) |
| **101** | L (0.999823) | L (0.999728) |
| **102** | A (0.999353) | A (0.999) |
| **103** | E (0.998666) | E (0.98408) |
| **104** | H (0.984462) | H (0.999627) |
| **105** | I (0.991805) | I (0.840342) |
| **106** | H (0.979793) | H (0.990368) |
| **107** | V (0.989116) | V (0.996634) |
| **108** | R (0.968539) | R (0.997839) |
| **109** | ***G (0.99667)*** | ***D (0.991238)*** |
| **110** | A (0.999353) | A (0.999) |
| **111** | ***N (0.99357)*** | ***R (0.988165)*** |
| **112** | L (0.999823) | L (0.999728) |
| **113** | E (0.987968) | E (0.998552) |
| **114** | N (0.999721) | N (0.999542) |
| **115** | G (0.999955) | G (0.999926) |
| **116** | L (0.999823) | L (0.999728) |
| **117** | L (0.999823) | L (0.999728) |
| **118** | Y (0.999936) | Y (0.999865) |
| **119** | I (0.998472) | I (0.997904) |
| **120** | D (0.999116) | D (0.998989) |
| **121** | L (0.999751) | L (0.999724) |
| **122** | E (0.999305) | E (0.999025) |
| **123** | R (0.999613) | R (0.999411) |
| **124** | V (0.770855) I (0.228497) | V (0.794577) I (0.204744) |
| **125** | V (0.904419) | V (0.990099) |
| **126** | P (0.99996) | P (0.999935) |

| | | |
|---|---|---|
| **127** | E (0.999305) | E (0.999025) |
| **128** | A (0.988944) | A (0.985575) |
| **129** | M (0.999359) | M (0.999806) |
| **130** | K (0.99958) | K (0.999428) |
| **131** | P (0.99996) | P (0.999935) |
| **132** | R (0.999613) | R (0.999411) |
| **133** | R (0.999523) | R (0.999404) |
| **134** | I (0.998482) | I (0.997912) |
| **135** | E (0.999209) | E (0.997485) |
| **136** | I (0.998482) | I (0.997912) |
| **137** | ***N (0.511055) K (0.449598)*** | ***L (0.988567)*** |
| **138** | ***-*** | ***K (0.619166) S (0.322748)*** |
