## supplementary file 4 - branch and branch - site model statistics for "Evolution towards simplicity in bacterial small heat shock protein system"

**Supplementary file 4A Branch model statistics for IbpA orthologs from *Enterobacterales*:** Models assumed either branch between nodes $AncA_0$ and $AncA_1$ or entire *Erwiniaceae* clade as foreground; NS – Not significant

| model | Np | lnL | ω estimates | LRT | |
|---|---|---|---|---|---|
| **M0** | **154** | **-11631.91** | **ω=0.04** | **N/A** | |
| **Two – Ratio, $A_0$-$A_1$ branch as foreground** | **155** | **-11618.80** | **$\omega_0$=0.04**<br>**$\omega_1$=999.00** | **LRT = 26.22** | **P<0.001** |
| **Two – Ratio, Erwiniaceae clade as foreground** | **155** | **-11630.36** | **$\omega_0$=0.04**<br>**$\omega_1$=0.04** | **LRT = 3.11** | **NS** |

**Suplementary file 4B Branch - site model statistics for IbpA orthologs from *Enterobacterales*:** Models assumed either branch between nodes $AncA_0$ and $AncA_1$ or entire *Erwiniaceae* clade as foreground; NS – Not significant

| model | | Np | lnL | site class | proportion | Background ω | foreground ω | LRT | |
|---|---|---|---|---|---|---|---|---|---|
| $A_0$-$A_1$ branch as foreground | A null | 156 | -11576.80 | 0 | 0.35 | 0.03 | 0.03 | N/A | |
| | | | | 1 | 0.01 | 1.00 | 1.00 | | |
| | | | | 2a | 0.63 | 0.03 | 0.40 | | |
| | | | | 2b | 0.01 | 1.00 | 0.40 | | |
| | A | 157 | -11565.20 | 0 | 0.88 | 0.03 | 0.03 | LRT = 23.20 | P<0,001 |
| | | | | 1 | 0.01 | 1.00 | 1.00 | | |
| | | | | 2a | 0.10 | 0.03 | 999.00 | | |
| | | | | 2b | 0.00 | 1.00 | 999.00 | | |
| *Erwiniaceae* clade as foreground | A null | 156 | -11523.90 | 0 | 0.90 | 0.03 | 0.03 | N/A | |
| | | | | 1 | 0.03 | 1.00 | 1.00 | | |
| | | | | 2a | 0.07 | 0.03 | 0.40 | | |
| | | | | 2b | 0.00 | 1.00 | 0.40 | | |
| | A | 157 | -11570.20 | 0 | 0.98 | 0.03 | 0.03 | LRT= - 92.51 | NS |
| | | | | 1 | 0.01 | 1.00 | 1.00 | | |
| | | | | 2a | 0.01 | 0.03 | 1.00 | | |
| | | | | 2b | 0.00 | 1.00 | 1.00 | | |
